# BrainTACO: An Explorable Multi-Scale Multi-Modal Brain Transcriptomic And Connectivity Data Resource

**DOI:** 10.1101/2023.04.18.537294

**Authors:** Florian Ganglberger, Markus Toepfer, Dominic Kargl, Julien Hernandez-Lallement, Nathan Lawless, Francesc Fernandez-Albert, Wulf Haubensak, Katja Bühler

**Affiliations:** Biomedical Image Informatics, VRVis Research Center, Vienna, Austria; Department of Neuronal Cell Biology, Vienna Medical University, Vienna, Austria; Global Computational Biology and Digital Sciences, Boehringer Ingelheim Pharma, Biberach an der Riss, Germany; Research Institute of Molecular Pathology (IMP), Vienna Biocenter (VBC), Vienna, Austria

**Keywords:** gene expression, brain connectivity, RNA sequencing, reference space, mapping

## Abstract

Exploring the relationships between genes, brain circuitry, and behaviour is accelerated by the joint analysis of a heterogeneous sets form 3D imaging data, anatomical data, and brain networks at varying scales, resolutions, and modalities. Hence, generating an integrated view, beyond the individual resources’ original purpose, requires the fusion of these data to a common space, and a visualization that bridges the gap across scales. However, despite ever expanding datasets, few platforms for integration and exploration of this heterogeneous data exist. To this end, we present the *BrainTACO* (Brain Transcriptomic And Connectivity Data) resource, a selection of heterogeneous, and multi-scale neurobiological data spatially mapped onto a common, hierarchical reference space, combined via a holistic data integration scheme. To access *BrainTACO*, we extended *BrainTrawler*, a web-based visual analytics framework for spatial neurobiological data, with comparative visualizations of multiple resources for gene expression dissection of brain networks with an unprecedented coverage. Using this platform, allows to straightforward explore and extract brain data for identifying potential genetic drivers of connectivity in both mice and humans that may contribute to the discovery of dysconnectivity phenotypes. Hence, *BrainTACO* reduces the need for time-consuming manual data aggregation often required for computational analyses in script based toolboxes, and supports neuroscientists by focusing on leveraging the data instead of preparing it.

## 1 Main

An increasing amount of evidence suggest that behaviours, and their impairments in psychiatric disorders, are better accounted by a multimodal data integration approach than when taking different neurobiological measures single handedly [1, 2]. Many insights into brains functional organization and neuronal mechanism were sparked by collecting and interpreting spatially organized histology, cellular composition, connectivity and activity data. For instance, the entry point for modern neuroscientific experimental workflows are brain regions (part of a specific neuronal circuit thought to be involved in a brain function or behaviour) whose gene expression and functional connectivity patterns are studied to understand the circuit dynamics underlying a behaviour. That information can then be used to identify targets in the brain, that could be modulated by psychoactive drugs, in cases of psychiatric symptomatology [3].Thus, integrating both functional connectivity and omics data modalities is instrumental in a better understanding of the biological underpinnings of behaviours and their deficits [4].

Recent advances in neuroimaging allowed big brain initiatives and consortia to create vast resources [5–7] of such data, which could be mined for additional and deeper insights. However, collecting these data from different sources for comparison and exploration leads to several challenges, as they are acquired in different systems, and can vary in resolution, anatomical scale, or sampling density. A mandatory first step is to map the data onto a common reference space, to ensure alignment (for imaging data) and annotation using the same brain region ontology [8]. The alternative approach, i.e., mapping the data to the smallest common denominator, such as major anatomical brain regions [9], one loses granularity and specificity, rendering the data potentially less representative.

Neuroscience studies that use a combination of omics, imaging, anatomical, and connectivity data often require extensive analytical workflows involving, mapping to a common reference space [8], manual data aggregation [9], and statistical analysis. This typically requires the expertise of a bioinformatician to find patterns that might relate to a given behaviour [9–16]. The term “big” refers to the amount (vast image collections, many datasets) and/or size (high resolution imaging/network data) of the data which is too extensive to analyse with traditional methods. Here, visual analytics tools bridge this gap by enabling neuroscientists to interactively browse vast data collections, visualize complex relationships, and link different types of data.

Many neuroscientific resources for transcriptomic data provide interactive, web-based visualizations for access and exploration. A comprehensive collection of such websites has been provided by Keil et al. [17]. While providing access to scientists without the need for advanced computational expertise, they are primarily suited for single datasets, i.e., they rarely provide workflows across multiple datasets and modalities. One notable exception is the SIIBRA-Explorer via EBRAINS [18] which combines structural connectivity (fibre tracts) [19] with microarray-based gene expression [20]. Another relevant tool, although not web-based, is BrainExplorer [21, 22], which enables the retrieval of structural connectivity from the Allen Mouse Brain Connectivity Atlas [23] in combination with in situ-hybridization data [24]. Nevertheless, in its current state both SIBBRA-Explorer and BrainExplorer are limited to one connectivity and one transcriptomic dataset each, and lack support for next-generation sequencing data.

In this paper, we present a holistic data integration scheme to map heterogeneous brain data across scales, spatial and anatomical resolutions, as well as sampling and acquisition types (Figure 1). *BrainTACO* (Brain Transcriptomic And Connectivity Data) is a resource that includes bulk and single-cell/nucleus RNA sequencing, in-situ hybridization, and microarray-based transcriptomics data, as well as structural and functional connectivity mapped onto common hierarchical reference spaces. To make *BrainTACO* accessible, we built onto previous work, *BrainTrawler* [25], a tool for visualizing volumetric, geometry, and connectivity data simultaneously in 3D rendering and 2D slice views, which can iteratively integrate additional heterogeneous datasets from the community and across species. We extended *BrainTrawler* to integrate, store and query datasets from various resources. Via a previously introduced data structure based on spatial indexing [26], it enables the automatic aggregation and interactive exploration in large-scale, high-resolution spatial connectivity [7, 23], and image collections of gene expression data [24] on different scales. We extended this data structure to integrate sample-based region-level datasets (i.e. sampled from a brain region), such as microarray gene expression data or count matrices from RNA sequencing. Here, it is possible to aggregate samples on individual dataset-level by user-defined regions of interest in real-time, so that different datasets can be compared on the same anatomical level, independent of their original resolution and scale.

**Fig. 1.**
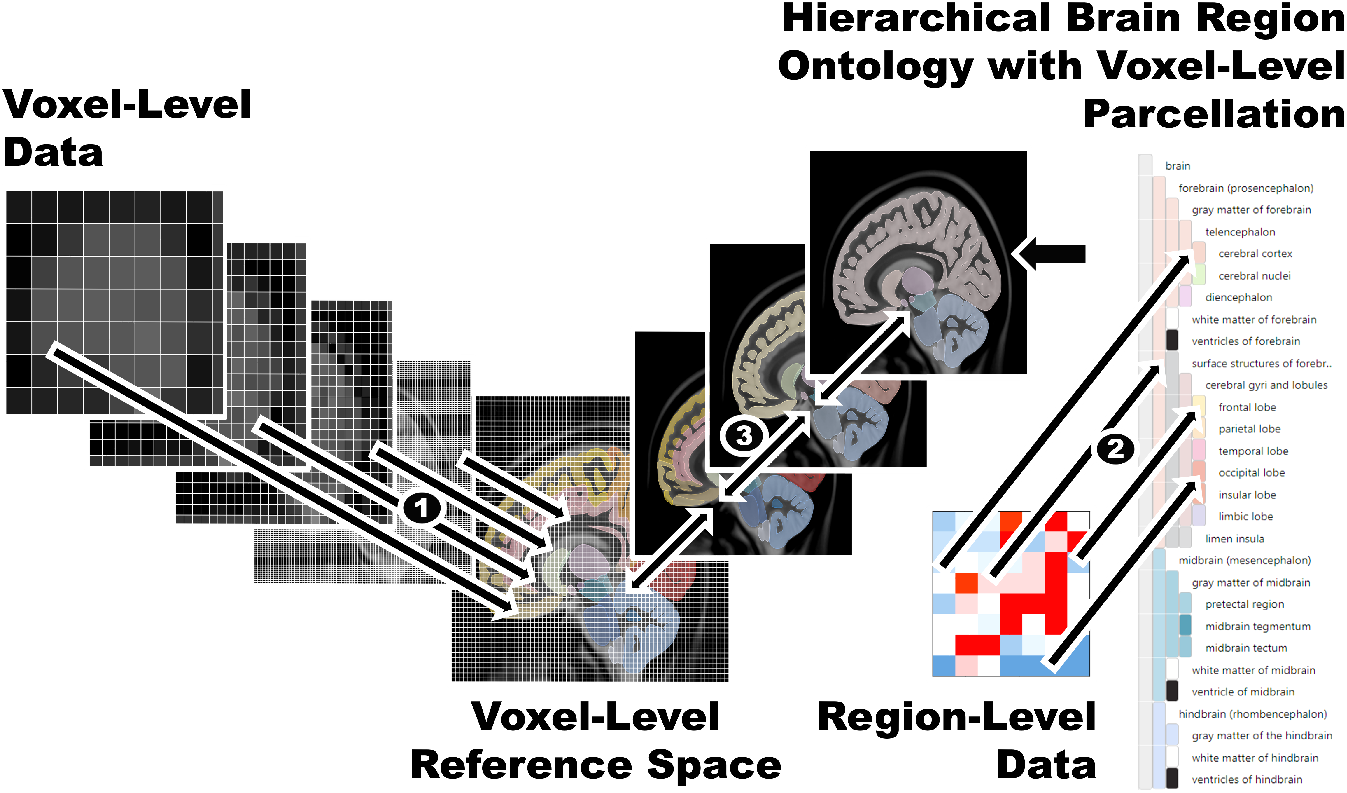
Mapping data of different resolution and scales to a common reference space. 1) Voxel-level data is mapped to the voxel-level reference space by image registration. 2). Region-level data (e.g. RNA-sequencing data) is mapped via a hierarchical brain region ontology with voxel-level parcellation to the reference space. 3) All data that have been mapped to the reference space can be either retrieved on the resolution of the reference space (data is up- or downsampled via nearest-neighbour interpolation), or on every other region level of the hierarchy.

Via a web interface, *BrainTACO* can be used to dissect brain connectivity interactively with a wealth of transcriptomic data, in a similar way as previously shown by Ganglberger et al. [25] for in-situ hybridization data only. To account for the increased number of datasets as well as the increased complexity of the datasets itself (e.g. samples from multiple cell types, developmental states etc.), we added additional comparative exploration. Here, we facilitated visualization techniques such as heatmaps, small multiples [27], and parallel coordinates to identify gene expression patterns across datasets and categorical information cell types, phenotypes, developmental stage) interactivity on arbitrary levels of anatomical detail. This enables neuroscientists a view on the data, tailored to their research focus, and without the need for programming knowledge.

This resource closes the gap in current interactive analytical tools by combining gene expression, structural, and functional relationships at the microscopic, mesoscopic, and macroscopic level. This is achieved by the following:

- A data hierarchical brain ontology-based integration scheme to access neurobiological, spatially mapped data across resolution, anatomical scale, or sampling density.
- A collection of publicly available gene expression (in-situ hybridization, microarray, bulk and single-cell/nucleus RNA sequencing) and connectivity (structural and functional resting-state) datasets covering major anatomical brain regions mapped onto common hierarchical reference spaces. The data’s original annotation is stored and made transparent (data provenance)
- An intuitive web interface for comparative visualization to access the *BrainTACO* resource in real-time without programming knowledge.

## 2 Results

### 2.1 Integrating multi-modal multi-scale resources

To create a resource of brain-wide gene expression and connectivity, we mapped heterogeneous neurobiological spatial datasets to common mouse [28] and human[29] reference spaces. We included a range of single-cell/nucleus RNA sequencing datasets (Figure 2) covering both species. While the datasets were representative of the whole mouse brain [30–36], the gaps in human data (e. g. Amygdala, Thalamus, Hypothalamus) [9, 34, 37, 38] were filled using bulk RNA sequencing datasets (Figure 2, GTEx and BrainSpan [39, 40]). The included datasest were selected to cover a diversity of meta information, such as morpho-electric cell types (patch sequencing [34]), age information (BrainSpan [40], Battacherjee at al. [33], Lee et al. [38], and the *STAB* datasets [9, 37, 41– 51]), or different treatment groups (Rossi et al. [32] and Battacherjee at al. [33]).]). To increase spatial resolution, we added in-situ hybridization data (200 micron voxel-level resolution) [24] and microarray gene expression data for 3702 biopsy sites [20], both already mapped to the reference spaces.

**Fig. 2.**
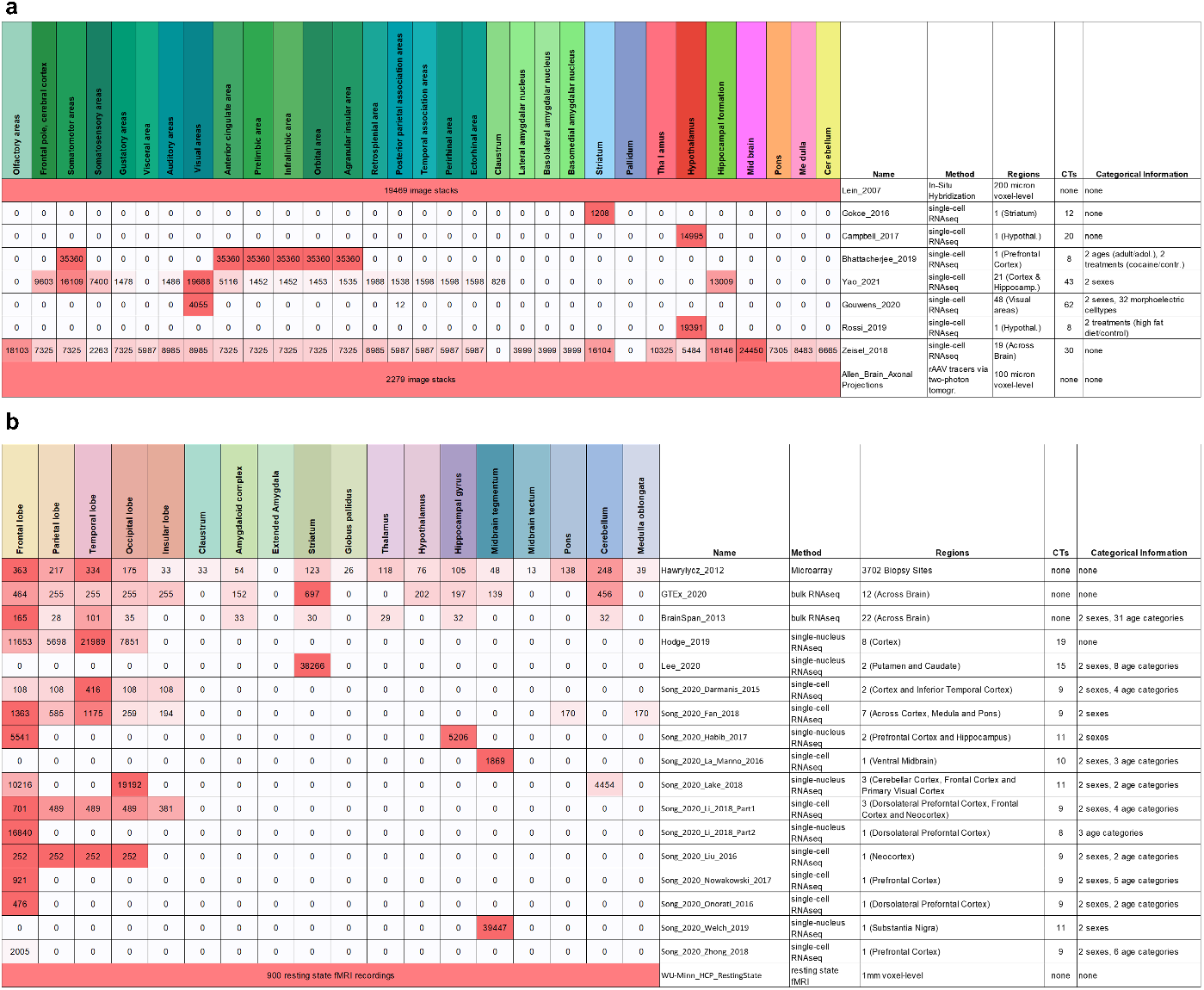
Dataset coverage over major anatomical brain regions. Numbers indicate the sample size/number of images of the datasets in the respective brain regions. Brain region colours represent the used hierarchical brain ontologies from the Allen Institute. a) Mouse datasets. b) Human datasets.

RNA sequencing datasets were leveraged using TPM (transcripts per million) for bulk RNA sequencing, and CPM (counts per million) for singlecell/nucleus RNA sequencing to ensure intra-dataset comparability of gene expression [52], at the expanse of inter-dataset comparability, which cannot be assumed due to technical biases [53] and different experimental conditions and/or sequencing protocols [52]. To circumvent this issue, two steps were taken. First, we limited the comparison to samples from adult subjects to avoid confounding due to varying developmental stages [9]. Second, inter-dataset comparability was assessed on rank level, i.e., whether the general order of genes by their TPM/CPM was consistent across datasets. Since similar brain regions were sampled in different datasets (e.g. Battacherjee at al. [33] and Yao et al. [35]), we computed the Spearman rank-correlation coefficient across datasets and modalities, which revealed consistent gene expression within specific cell types across datasets (Figure 3, code and additional information in Supplementary Data 2 and Supplementary Table 1). For details about the mapping of the datasets, as well as the preprocessing and normalization, see the *Methods* (Section 4.2 and 4.1).

**Fig. 3.**
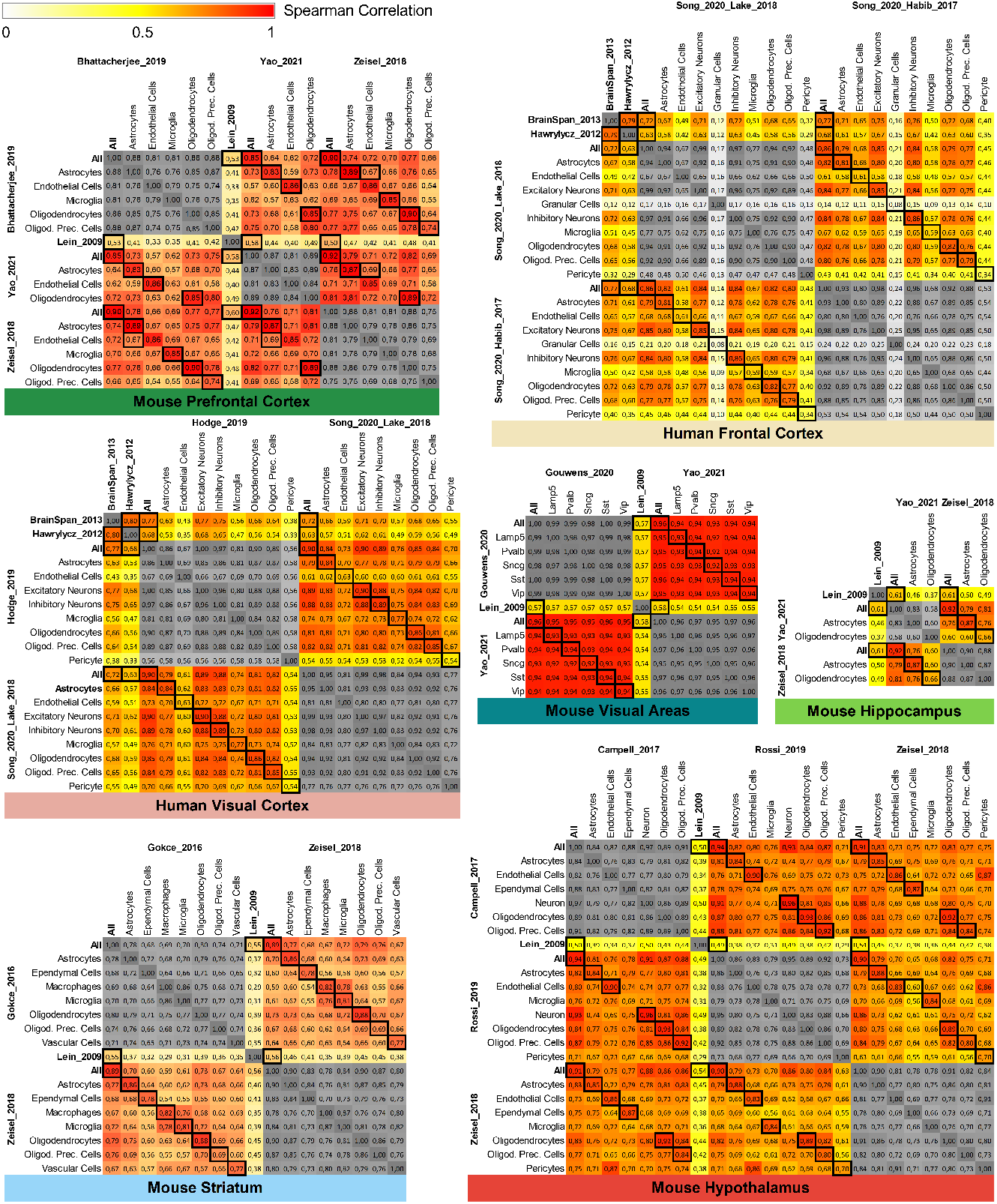
Heatmaps showing the (Spearman rank-based) correlation over all genes of mouse and human datasets that cover the same brain regions and cell types. Black boxes mark correlations of the same cell types in different datasets. On average, their correlation is higher than for not-matching cell types, which indicates that (the ranking of) gene expression is consistent across datasets (one-sided Wilcoxon test, all p-values≤ 0.05, except mouse visual areas). Neuronal subtypes in mouse visual areas were already so similar within datasets (all correlation≥ 0.95), that there was no significant difference across cell types.

### 2.2 Mapping to a common reference space

The joint exploration of spatial datasets from different resources requires the data to be aligned to a common space [8]. This space acts as a reference, so that spatial locations, such as coordinates or brain region annotations, have the same meaning across datasets. In neuroscience, commonly used reference spaces are typically defined by an anatomical reference template [28, 29], a structural image that has been combined (e.g. via image registration) to a structural representation of the brain for a group of specimen or a species.

Imaging data, i.e., data that is divided by a 2D/3D grid into pixels/voxels that represent measurements at their respective positions, can then be aligned onto a template by image registration. This involves image transformation and warping to establish voxel-level correspondence (Figure 1). As at templates we used the Allen Mouse Brain Coordinate Framework [28] for the mouse brain (0.1 mm resolution) and the ICBM 152 MNI space template [29] (1 mm resolution) for their widespread use and availability [54]. In principle, there is no limitation to specific templates.

For spatial data that is not derived from imaging, i. e., measurements that have only been generated for specific brain regions, a different approach is needed. Here, we utilised hierarchical ontologies of brain regions, a formal representation of knowledge about the species-specific brain anatomy [55] (Figure 1), i.e., which brain regions it consists of and how these brain regions are subdivided (hierarchically). The Allen Institute provides ontologies for both mouse and human [28, 56, 57] that include a mapping onto our respect reference spaces, i.e., which coordinate of the reference space belongs to which brain region in the ontology. Since datasets are not necessarily annotated with the same ontology and on the same hierarchical level, they cannot be compared across anatomical scales and resolution directly. Hence, we matched brain region annotations to the corresponding brain regions in the ontology. An outline of this process is shown in Figure 1, details can be found in the *Methods*, Section 4.2. Note that these mappings are made explicit in our resource’s user interface to ensure transparency, and as a consequence, allow for quality control.

The distributed nature of brain functions across brain networks and gene sets required adequate exploration of spatial gene expression in the context of brain structural and/or functional connectivity. We therefore integrated high-resolution imaging data of structural connectivity (for mouse) [23] and resting-state functional connectivity (for human) [7]. Structural connectivity describes how brain areas are physically connected via axonal projections, and was originally imaged on a 100 micron resolution [23]. The human resting-state functional connectivity describes brain regions that are linked by correlated activity. This data originated from the WU-Minn Human Connectome Project [7]and represents the group-average dense, voxel-level correlation of the resting-state BOLD-signal of 820 subjects.

### 2.3 Interactive access and exploration

*BrainTrawler* was one of the first iterations of an interactive, web-based framework visual analytics framework [25]. Originally, it was designed to explore large-scale brain connectivity data, such as structural connectivity [23] and to dissect these connections on gene expression level in the mouse brain. This was achieved by providing a visual analytics workflow to identify which genes are expressed in either the source or target regions of these connections, by including spatially mapped gene expression data of 20.000 genes of the Allen Mouse Brain Atlas [24]. Interactivity was achieved by facilitating spatial indexing on volumetric images [58] for the spatially mapped gene expression data, as well as a data structure for real-time aggregation of connectivity data with billions of connections [26].

We here build on this effort to handle large-scale transcriptomic datasets for mouse and human, to not only showing where genes are expressed, but also how expression differs between cell types and developmental or physiological conditions. Here, we build a spatial database of RNA sequencing and microarray-based gene expression datasets, including the datasets described in the previous section. This spatial database utilizes spatial indexing for aggregating gene expression of datasets in real time, that were aligned to brain regions/voxels of the reference space. To this end, datasets including their meta data (e.g. cell type annotations, age, phenotype, etc.) were sorted based on their spatial location in the brain (see *Methods*, Section 4.4 for details).

The exploration of gene expression related to brain connections works in an analogous manner as previously presented in Ganglberger et al. [25], see Figure 4: First, the user defines a volume of interest (*VOI*), which can be either an arbitrary manually defined area, or a brain region (Figure 4a, yellow area). For this *VOI*, a gene expression query can be performed, which computes the mean expression of all datasets that have been aligned to the reference space within the *VOI* or a user-defined filter, i.e., selected meta properties data such as certain cell types, phenotypes, and others (details in the *Methods*, Section 4.2).

**Fig. 4.**
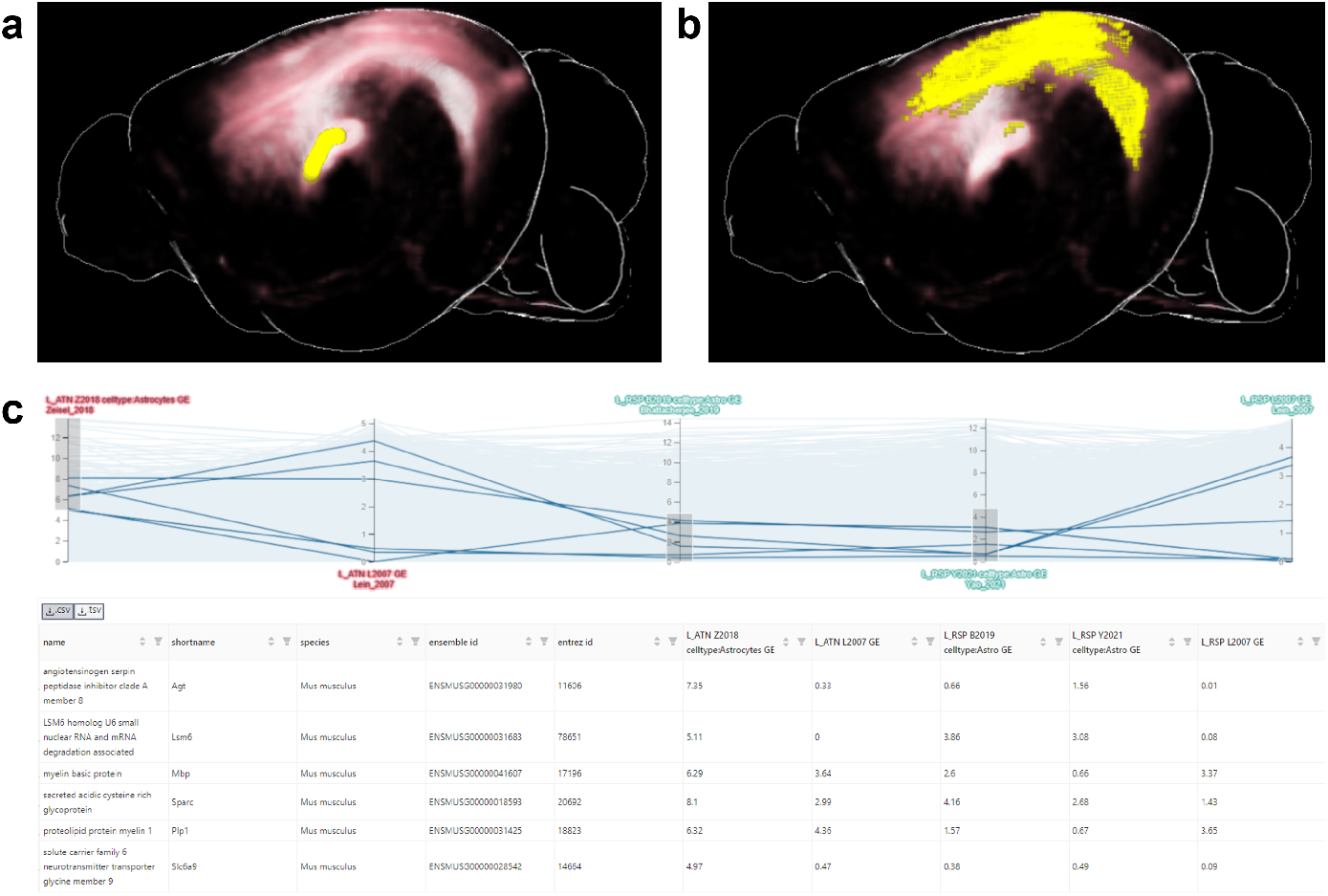
Exemplary gene expression dissection of a structural connection. a) Outgoing structural connections (red) from a user-selected part of the Thalamus (yellow, representing the source VOI of the connection). b) Target VOI (yellow) of the structural connections (red). c) Exemplary gene expression in a parallel coordinates system from the source VOI (red axes, one for astrocytes in the Zeisel 2018 [36] dataset, one for gene expression in the Lein 2007 dataset) and target VOI (green axes, gene expression in the Lein 2007 dataset, Astrocytes in the Battacherjee 2019 [33] and Yao 2020 datasets [35]). Every horizontal line represents a gene, their location on the axes their level of gene expression. A subset of genes with low astrocytes expression in the source VOI and high Astrocytes expression in the target VOI was selected as an example.

The result of such a query is a list of genes with the aggregated gene expression. Figure 4c shows how multiple queries results can then be compared in a parallel coordinate system, which allows filtering multiple gene lists by their gene expression. Each axis in the figure represents the result of a gene expression query, and, as a consequence the level of gene expression in the query regions. Each horizontal line represents a gene. A selection/filtering of genes (shown in the table in the lower part of Figure 4c) with specific gene expression patterns can be made drawing brushes on an axis. Since queries of different *VOI* s can be compared, one can use this on the source and target areas from connectivity data for gene expression dissection. Figure 4a and b show the aggregated outgoing structural connectivity of the *VOI* in red. While the yellow *VOI* in Figure 4a represents the source, the yellow area in Figure 4b represents the (strongest) targets of the aggregated connections. A comparison of the gene expression of source and target *VOI* can be seen in Figure 4c. Here, the axes labeled in red are results of gene expression queries at the source *VOI*, green ones at the target *VOI*, performed for different exemplarily selected datasets and cell types.

The increasing dataset number and complexity (e.g. samples from multiple cell types, developmental states, etc.) makes it necessary to perform large amount of expression queries to cover all available information for genes of interest. Hence, we extended *BrainTrawler’s*capability to visualize gene expression of multiple resources jointly by developing a lightweight interface (*BrainTrawler LITE*). *BrainTrawler LITE’s* basic user interface element is a heatmap of the dataset coverage (Figure 5a). Here, each heatmap tile represents the sample size/image number distribution of a certain dataset (rows) for a certain brain region (columns), similar to Figure 2. By clicking on heatmap tile, these data can be selected for further investigation: Either on a gene set level, by entering a list of genes (Figure 5b), or on a genome-wide level (Figure 5c), analogously to a gene expression query. Results can be exported as images or as comma/tab-separated files for later use or for sharing. For more details see Methods, Section 4.3.

**Fig. 5.**
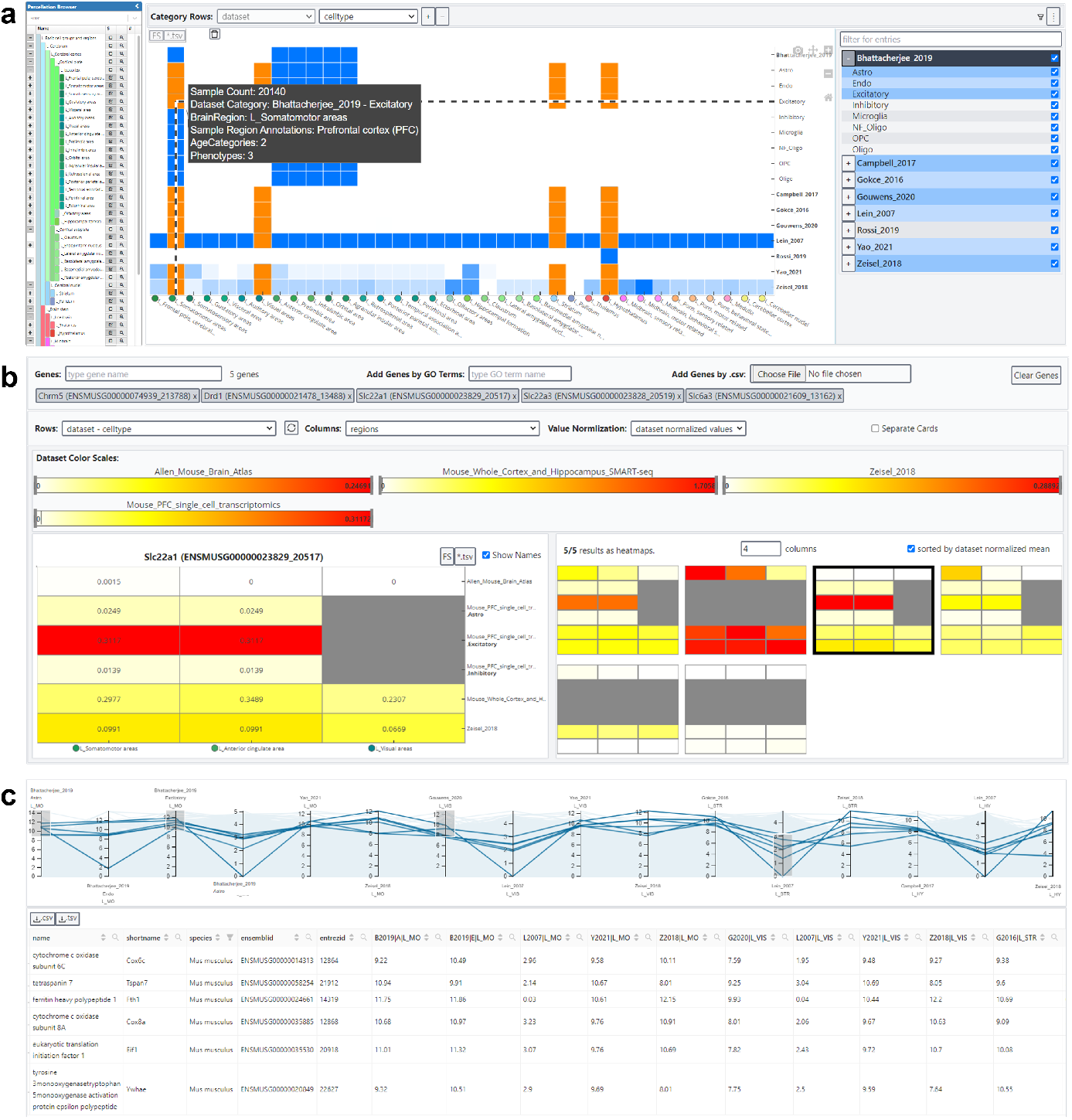
BrainTrawler LITE interface for comparative visualization of gene expression across datasets. a) The dataset coverage heatmap shows the distribution of sample size/image number across brain regions (columns) and datasets (rows), subdivided by meta data attributes such as cell types, phenotypes, etc. Brain regions and meta data categories can be adapted via tree-like UI elements on the sides, the tooltip shows the exact composition (sample/image count, meta data categories, etc.) of the respective heatmap tile. Orange tiles shows a selection of data for gene expression visualization (in b and c). b) Gene expression heatmaps of five selected genes. Rows and columns represent the selected (orange) tiles in the dataset coverage heatmap (in a). Colour scales are separated per dataset (between 0 and the maximum value shown for all selected genes). The right side shows the expression of genes for each dataset separately as small multiples, the left side shows one selected gene with more details (labels, values etc.). Grey tiles are missing data. c) Parallel coordinates system showing the gene expression of all genes in the selected dataset on axes, each representing the average expression of genes (blue lines) of the samples/images of each selected (orange) tile in the dataset coverage (in a). Via drawing brushes on the axes, the genes in the parallel coordinates system can be filtered. Filtered genes are shown below in a table.

### 2.4 Relating gene expression and connectivity across species uncovers genes and mechanisms for human functional connectivity

The functional (FC) and structural connectome (SC) of the brain is viewed as a major determinant of cognitive function across species. Altered connection topology and intensity of brain areas are common correlates of psychiatric conditions such as the autism spectrum and schizophrenia, suggesting that dysconnectivity might lie at the core of these conditions [59–61]. Alongside, GWAS studies have discovered genetic loci and polymorphisms associated with these psychiatric conditions, suggesting that the relationship between the connectivity of a given brain area and its gene expression likely harbours valuable information on wiring principles of the brain [62, 63]. To bridge these domains, the combination of connectomic and gene expression data is a promising approach to discover genetic etiologies and emerging mechanisms that drive regional efferent and afferent connectivity underlying connectopathies, conditions associated with aberrant brain connection topology [64].

Thus, the increasing abundance of (cell type-specific) gene expression and connectivity data in the mouse is a promising avenue to discover genetic susceptibilities and mechanisms affecting the connectome, with great translational potential. Thus, identifying genetic drivers of FC in humans is of great interest, as it may provide entry points into therapeutic interventions to ameliorate disease burden.

In the context of psychiatry, the insular cortex is of special interest, as a core hub in regulating large scale brain networks in humans [65] and rodents [66], while involved in interoceptive, cognitive and affective processes [67, 68]. Notably, insular functional dysconnectivity is a signature of common psychiatric conditions [69, 70]. We segmented the insula into its agranular and granular portions, as they are anatomically and functionally distinct [71]. While the posterior granular insula (GI) is a primary sensory area with rich afferents for interoceptive information, the anterior agranular insula (AI) represents an associative area with increasing multimodal integration [67]. Therefore, this system is an ideal model to discover novel genetic factors shaping the connectivity of cortical areas of distinct architecture (agranular vs granular insula), in a highly relevant translational setting.

First, to allow for optimal cross-species inferences, we selected consensus areas between the rodent and human brain, covering 10 major subcortical areas (Supplementary Table 2). Next, source and target connectivity data with these areas was sampled for the AI (combined ”L_Agranular insular area, dorsal part” and ”L_Agranular insular area, ventral part”) and GI (”L_Visceral area”) (Supplementary Figure 1, left). Because the human ABA does not discern by granularity, human FC data for AI and GI with the consensus areas was sampled by brushing agranular and granular areas of the short and long insular gyri (Supplementary Figure 1, right, according to [72]. Within species analysis shows a correlation between source and target SC in the mouse GI, but not in the AI (Figure 6a). Overall, AI and GI connectivity is correlated in rodents and humans (Supplementary Figure 1a), although to different extents between sources and targets. Interestingly, human FC is not significantly correlated to mouse SC, suggesting relevant functional differences between species and/or connection modality (Figure 6b).

**Fig. 6.**
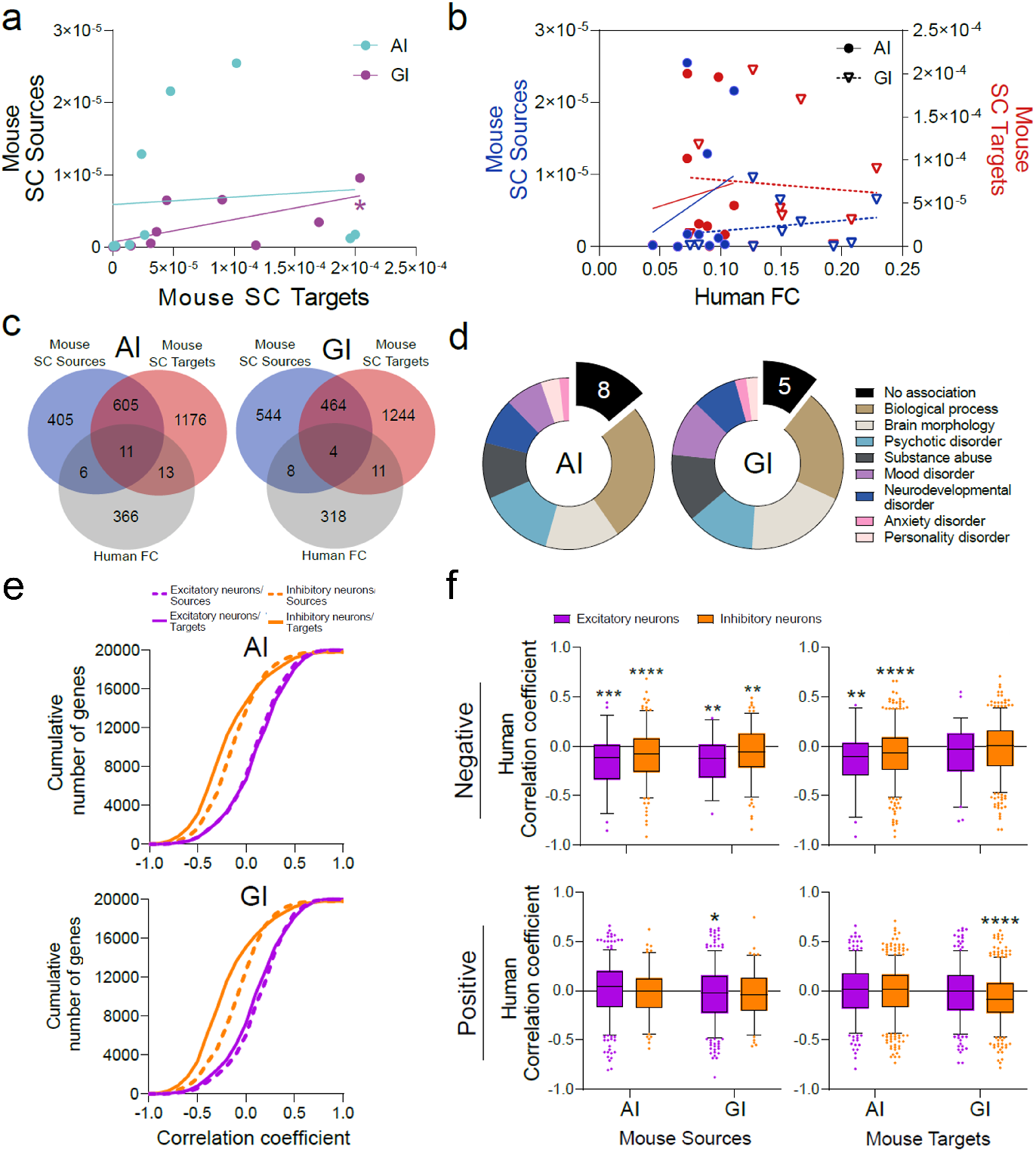
Relating regional gene expression with connectivity across species identifies potential connectivity driver genes, where cell type-specific analysis reveals a conserved inhibitory mechanism across species and connection modality. a) Mouse source and target connectivity is significantly correlated for GI (Spearman r =0.65, p-value = 0.04), but not for AI (r = 0.08, p-value = 0.82). b) Mouse SC source and target connectivity are not significantly correlated to Human FC (Spearman; AI: source r = 0.22, p-value =0.54, target r = 0.22, p-value = 0.54; GI: source r = 0.22, p-value = 0.54, target r = −0.05, p-value = 0.89). c) Overlap of genes with expression significantly correlated to connectivity across species (Mouse SC source/target and Human FC). d) Brain-related associations of overlapping genes in c (total of 30 genes for AI, 23 genes for GI; see Table 4 for summary). e) Cumulative distribution of correlation coefficients resulting from the correlation of cell type specific gene expression and Mouse SC (sources/targets) across excitatory and inhibitory cell types for AI and GI. Kolmogorov-Smirnoff tests revealed significant differences for AI and GI (p-value ≤ 0.0001) between excitatory and inhibitory cell types for within source and target connectivity, respectively. In addition, significant differences (p-value ≤ 0.0001) between source and target connectivity within excitatory and inhibitory neurons, respectively, were found for both AI and GI. f) Selection of human correlation coefficients with significantly correlated homologs in the mouse, based on direction of correlation in the mouse dataset (top: negative, bottom: positive), mouse SC type (left: Sources, right: Targets) and cell type of origin (color) and connected region (AI, GI). Significance was tested by one-sample t-test against zero (chance level). ** p-value ≤ 0.01, *** p-value ≤ 0.001, **** p-value ≤ 0.0001.

To assess the relationship between gene expression and connectivity we extracted expression data of major excitatory and inhibitory cell types of the 10 subcortical consensus areas for which expression data is available from Zeisel 2018 [36] (Mouse) and Hawrylycz 2012. [20] (Human) (Supplementary Table 2). These were correlated with FC and SC within humans and mouse, respectively. This identified for AI 1027/2252 and for GI 1020/1723 significantly (anti-)correlated genes with mouse source/target connectivity, respectively. For human data we found 330 and 286 significantly (anti-)correlated genes with FC for AI and GI, respectively (Supplementary Table 3). To identify potential basic driver genes for insular connectivity (i.e. those conserved across the available mouse structural and human functional connectivity data), we determined the overlap between human and mouse datasets. This resulted in a total of 30 genes for AI (Figure 6c, left) and 23 genes for GI across mouse sources and targets (Figure 6c, right; see Supplementary Figure 2 for source- and target-specific analysis and Supplementary Table 4). Association analysis for brain-related categories on these genes in *Opentargets* [73] suggests that they are involved in processes, relevant to psychiatric conditions (Figure 6d). In this context, our workflow recovers several genes previously associated with autism (AI: 3 genes; GI: 3 genes) and schizophrenia (AI: 8 genes; GI: 6 genes) (see Supplementary Table 5). Among the positively correlated we find attractin-like 1 (ATRNL1) specifically for AI, a gene previously found to be mutated in a human patient diagnosed with autism [74]. The estrogen receptor 2 (ESR2) is among few genes with a link to schizophrenia, to have switched from strongly positive correlation with AI (sources and targets) and GI connectivity (targets only) in mouse to a strong negative correlation in humans (Supplementary Table 5). However, we find several genes previously not linked to brain-related disorders, potentially identifying novel genetic factors that contribute to dysconnectivity phenotypes.

It is established that brain oscillations are governed by recurrent inhibitory networks [75], suggesting that genes expressed in inhibitory neurons might drive FC. To address this systematically, we harnessed the available cell type specific gene expression data in the mouse as an entry point. We extracted gene expression data of excitatory and inhibitory cell types from the 10 consensus areas in the mouse using “Cell type specificity” query in the mouse (Supplementary Table 2). This approach should emphasize cell-type specific genes and thus enhance contrast between cell types. We next correlated this cell-type specific gene expression data to the mouse SC of AI and GI (Sources and Targets) to the consensus areas. Indeed, the majority of significantly correlated genes are more specific to inhibitory neurons (Figure 6e). Interestingly, we noted that the majority of these genes are negatively correlated, suggesting inhibitory mechanisms might shape mouse SC.

We then tested, whether this principle is conserved across species and connection modalities. First, human correlation coefficients were selected for by significantly correlated homologs in the mouse dataset, maintaining source/target identity and cell type of origin and direction of correlation. This way, a conserved direction of correlation between species might uncover shared mechanisms. These human correlation coefficients were then tested for a deviation from zero (chance level). Indeed, genes negatively correlating with SC in the mouse statistically preserve the negative correlation to human FC. This is most dominant for genes negatively correlating with mouse source connectivity (Figure 6f, top left), and specific to AI for genes from mouse target connectivity (Figure 6f, top right). Interestingly, this pattern is absent when using mouse SC positively correlated genes, suggesting specificity for a conserved inhibitory mechanism (Figure 6f, bottom). Among the strongest hits we found is TNF superfamily member 12 member C (TNFSF12), a gene dysregulated in patients diagnosed with various psychiatric disorders [76–78].

In summary, despite the fact that the correlation of human FC and mouse SC does not reach significance (Figure 6b), this workflow uncovered a preserved relationship between gene expression and connectivity across species and connection modality. This may further allow to mine for (cell type-specific) genes and mechanisms driving FC in humans.

## 3 Discussion

Our platform has several implications for both basic and biomedical research. We created a discovery framework that utilizes data from current popular large-scale genetic and brain network initiatives to rapidly screen for neural circuitry underlying specific brain functions, behaviours, or psychiatric symptoms at comparably low computing costs. The computational screening complements, and may direct subsequent, circuit-genetic experiments such as electrophysiology, opto-, and pharmacogenetics. When performed at large scale with behaviour-specific genes, our approach has the potential to refine the functional organization of the brain beyond simple anatomical domains. Importantly, our methodology for generating and exploiting the resource could be applied to other model organisms for which spatially mapped gene expression, network, and genetic information is, or will be, available, for example fruit flies or zebrafish. Using this platform, we showcase this by uncovering a preserved relationship between gene expression and connectivity across species and connection modality. We explored this in excitatory and inhibitory cell types, and identified several genes that were previously not linked to brain-related disorders, potentially identifying novel cell type-specific factors that contribute to dysconnectivity phenotypes.

Inevitably, *BrainTACO* has some limitations. First of all, in its current state it does not cover the full brain on a single-cell/nucleus level. To provide a resource as versatile as possible, we focused on covering the brain at least on the level of most major anatomical brain regions, so that neuroscientists will likely find data related to their research focus. This was not entirely possible for human datasets (e. g. Amygdala, Thalamus, Hypothalamus) due to a lack of studies covering these areas. In principle, *BrainTACO* is not static, and can be extended with new datasets. Future studies, as well as further improvements of technologies such as spatial transcriptomics will help to close this gap.

Another limitation is that it is in general not reasonable to compare (absolute) gene expression values, even normalized ones (e.g. TPM, CPM) across datasets [52, 53]. We circumvent this issue by providing dataset-specific colour scales for gene expression heatmaps, and advise to compare absolute gene expression only relative to other genes within the same dataset. As an alternative, we provide gene ranks for gene expression queries, i.e., their relative position in a list of genes sorted by their expression. Furthermore, the computed mean expression, region and cell type specificity or enrichment scores do not include information about the spread of the data, which is typical for traditional analytical approaches, such as t-SNE plots. Hence, it is not known how homogenous the expression is across the selected datasets, for example for a certain cell type. We knowingly limited analytical power for the sake of simplicity, cornerstone of *BrainTrawler*, whose mission is to enable computationally agnostic neuroscientists to run complex analysis. Nevertheless, an integration of more sophisticated, in-depth analyses might be considered for future releases.

The user interface provides sufficient visualization for the scope of our case studies, i.e. the number of queries, datasets and genes did not exceed *BrainTrawler’s* capabilities. In general, there is no limit regarding how many genes, datasets or query results can be visualized, but outside typical analytical workflows, such as our case study, there are some issues regarding the user interface: The parallel coordinates system does not scale well to more than twenty queries, for there is not enough space in a typical browser window. The same is true for visualizing the expression of hundreds of genes in *BrainTrawler LITE’s* gene expression heatmaps, for you simply loosing the overview when scrolling is needed.

Overall, the integration of heterogeneous gene expression and connectivity data from mouse and human into *BrainTrawler* is a powerful resource for hypothesis building in the field of behavioural/functional neuroscience and for drug target identification. Its coverage of RNA sequencing data, especially on a single-cell/nucleus level for the majority of brain regions, only limited by the public availability of the data, enhances *BrainTrawler’s* capabilities of investigating molecular mechanisms. By making the *BrainTACO* resource available via visual analytics workflows in the web, we enable quick access without manual data aggregation via scripting, and consequently without the expertise of a bioinformatician. Future integration of novel spatial datatypes, such as spatial transcriptomics, has the potential to make this resource even more versatile.

## 4 Methods

### 4.1 Data Preprocessing and Normalization

- Single**-Cell/Nucleus RNA Sequencing Data:** We integrated 21 singlecell/nucleus RNA sequencing datasets in total, 7 mouse datasets, and 14 human. Out of the 14 human datasets, 12 are from the Song et al. [9] meta dataset STAB available from http://stab.comp-sysbio.org. STAB consists of 13 datasets [37, 41–51], for which we omitted the dataset by Hodge et al. [37] and downloaded the data from Hodge et al.’s original resource (http://celltypes.brain-map.org/rnaseq), since it had been extended by several brain regions after STAB’s submission (primary motor cortex, primary somatosensory cortex, and primary auditory cortex). The other singlecell/nucleus RNA sequencing datasets were obtained by the information in the data availability statements in their referenced publication. *STAB* datasets were not further preprocessed, filtered and normalized, since this has been already done consistent across its 12 datasets [9]. The other single-cell/nucleus RNA sequencing datasets were pre-processed with the Seurat (v4.1.0) R package [79] to remove batch effects. Low-quality cells, empty droplets, and doublets were removed with low (less then 50) or high (more than 5000) unique gene counts, or if their unique gene counts were outliers (lower or higher five times the median absolute deviation from the median). Final cell counts can be seen in Table 1 (Filtered Cell Counts). Note, that most available datasets were already filtered by similar criteria, which explains the similar or close original and filtered cell counts. Cells that could not be matched to brain regions were removed, which was only the case for samples from the medial ganglionic eminence (transient structure in the developmental brain) in the Nowakowski et al. [41] data from *STAB*. Genes without cell counts across the datasets where removed, since they do not show biological variability. Genes were then matched by gene symbol, ensemblid or entrezid to *BrainTrawler’s* gene database, which was obtained via the *Genome wide annotation* for mouse [80] and human [81] via the bioconductor package. The amount of matches can be seen in Table 1. For each gene, expression levels were normalized by computing CPM (counts per million) to ensure intra-dataset comparability of gene expression [52]. For better readability/interpretability, CPM was log2 normalized (using an offset of 1 to account for zeros).
- **Bulk RNA-Sequencing Data:** To fill gaps in subcortical singlecell/nucleus RNA sequencing availability for the human, we integrated two bulk RNA sequencing datasets from the GTEx and BrainSpan consortia [39, 40]. GTEx data was downloaded from the GTex portal (https://gtexportal.org) in the version 8 as gene TPM (transcripts per million). BrainSpan data was obtained from the BrainSpan portal (https://www.brainspan.org/) as normalized RPKM (reads per kilobase of transcript) expression values, and converted to TPM according to Zhao et al. [52]. log2 normalization (using an offset of 1 to account for zeros) was applied to both datasets similar to single-cell/nucleus RNA sequencing data.
- **Microarray Gene Expression Data:** Microarray gene expression data was retrieved from the Allen Human Brain Atlas by Hawrylycz et al. [20] via the Allen Brain Atlas API (https://api.brain-map.org/). This data assemble gene expression from 3702 samples of six donors, labelled with their according brain region in the ontology provided by the Allen Institute [56], which ensures equivalent scaling across donors. We normalized gene expression values based on an outlier-robust sigmoid function, before rescaling the normalized values to a unit interval (0-1), as suggested by Arnatkevĭciūtė et al. [82].
- **In-situ Hybridization Data:** Whole-brain gene expression in situ hybridization data was retrieved from the Allen Brain Atlas API (https://api.brain-map.org/) as volumetric images for 19479 genes. To create these volumetric images, the Allen Brain Atlas divides the in situ hibridization slice images on cellular resolution into a 200 micron resolution grid. For each grid division, expression energy was computed, i.e. the sum of the expression intensity of all pixels within the division, divided by the sum of pixels within the division [83]. The expression energy for each grid division represent then a 200 micron resolution volumetric images. To make this data available in *BrainTrawler*, we log2 normalized the data and encoded them as 8 bit volumes, with a size of 155KB each (*∼* 3*GB* in total).
- **Structural Connectivity Data:** Structural connectivity was generated similar to previous publications [25, 26, 84]. Here, the connectivity was retrieved from the Allen Brain Atlas API (https://api.brain-map.org/) as volumetric images, showing structural connectivity of 2173 injection sites to their target sites [23]. These 2173 images were generated on a 100 micron resolution by labelled rAAV tracers via serial two-photon tomogagraphy [23]. For each image, the injection site is given by coordinates in the reference space defined by the Allen Mouse Brain Coordinate Framework [28], and an injection volume, depicting the volume around the injection site affected by the tracer. Hence, the connectivity for an injection site is defined by the all voxels within its injection volume. For every voxel in the reference space, we took the connectivity from the covering injection volume. If a voxel was covered by multiple injection volumes, i.e., and therefore by multiple injection sites, we combined them by taking the maximum connectivity for each target. To compensate for low count of injection sites on the left hemisphere, we mirrored the connectivity, effectively inflating the original 2173 injection sites to the double, i.e., 4346. To minimize the amount of false positive connections, the data was thresholded by values *<* 10*^−^*^45^ according to Oh et al. [23], Extended Data Figure. The result was a dense *∼* 67500 *×* 500000 structural connectome (*∼* 67500 source voxel covering injection volumes with *∼* 500000 target voxels within the mouse brain), with *∼* 90*GB* stored in a csv format.
- **Resting-State** Functional **Connectivity Data:** Resting-state functional connectivity data was downloaded from the WU-Minn Human Connectome Project [7] via the CONNECTOMEdb (https://db.humanconnectome. org/). The data was available as average functional connectivity matrix of 820 subjects, given as dense *∼* 90000*×*90000 functional connectome in “gray-ordinate” space [7], where a grayordinate is either a voxel (subcortical gray matter) or a surface vertex (cerebral cortex). To transform this matrix into the ICBM 152 MNI space [29] (1mm resolution), we used the Connectome Workbench platform [7] to retrieve the closest grayordinates (within 1mm) to every voxel in ICBM 152 MNI space. By computing the average connectivity of all close grayordiantes for each voxel, we were able to create a dense *∼* 87000 *×* 87000 functional connectome, with *∼* 45*GB* stored as csv.

**Table 1.**
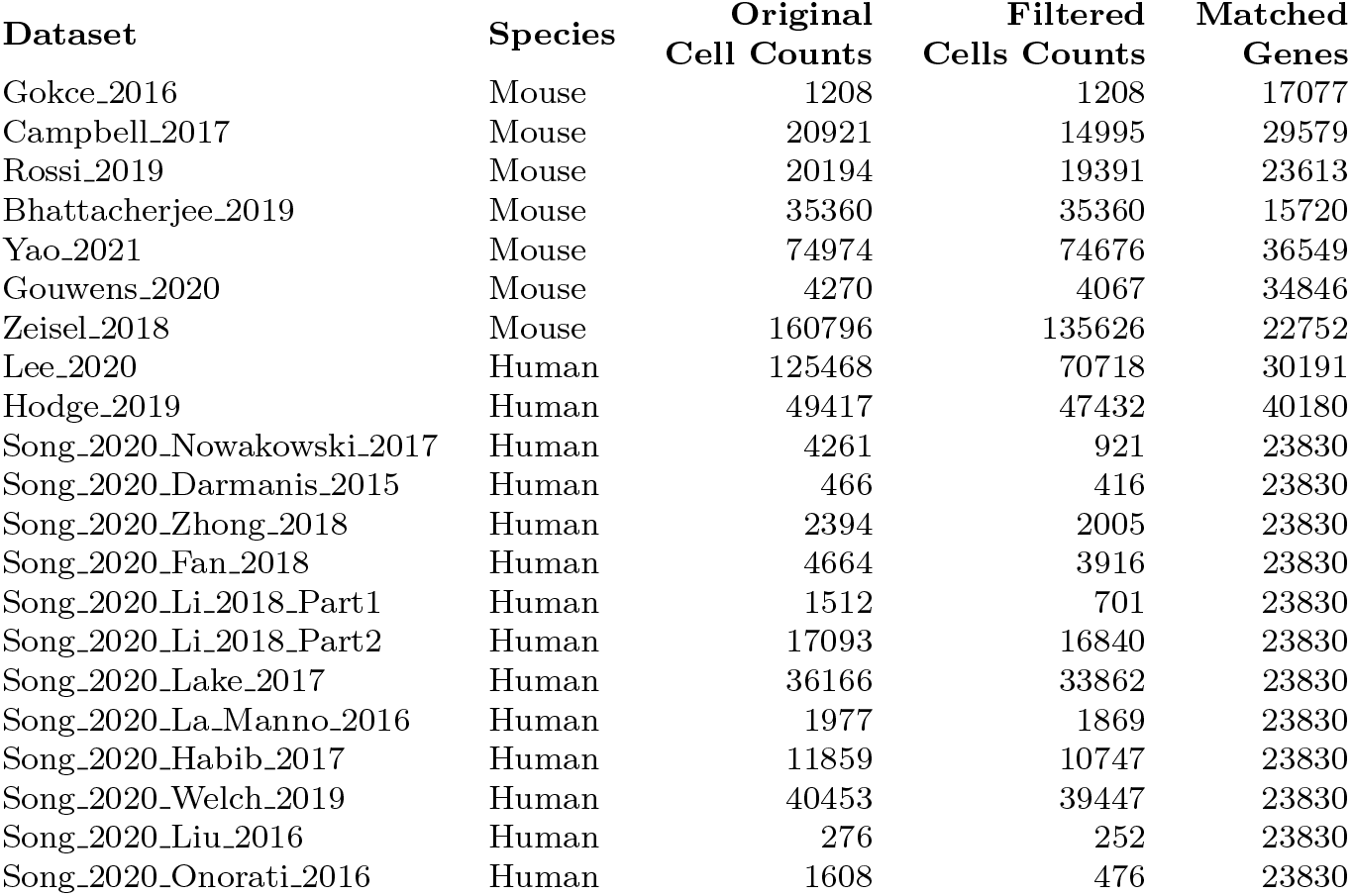
Original Cell Counts of the retrieved datasets, filtered cell counts after preprocessing, and genes matched to BrainTrawler’s gene database

### 4.2 Data Mapping and Querying

Imaging data that is shown in this paper, namely in-situ hybridization data [24], axonal projection connectivity [23] and resting state functional connectivity [7] was already aligned to the reference spaces used for this resource [28, 56, 57]. Novel datasets could be aligned via tools such as the QUINT workflow [85] or the ANTS frame work [86]. Data that does not meet the resolution of the reference space, is up- or downsampled via nearest-neighbour interpolation. This was the case for the in-situ hybridization data, which has a lower resolution (200 microns) than the Allen Mouse Brain Coordinate Framework [28] (100 micron resolution).

Region-level data, such as microarray gene expression data and RNA sequencing data are typically encoding gene expression as count matrix [87], depicting the frequency of gene transcripts for samples. For dataset included in this study, these samples originatee from brain regions. We mapped these brain regions to the corresponding brain regions of our reference ontology (Allen Brain Institute Atlases) manually based on the region name and description in the dataset’s reference publication. To ensure transparency, and hence quality control, the detailed mappings are available in the resource’s user interface (*Browse Database*, then select a dataset to see details such as the dataset’s mapping), and in the supplemental material.

The process of mapping region-level data to, and retrieving it from a reference space is outlined exemplarily in Figure 7, code for the mapping can be found in Supplementary Data 1. In this example, these data are samples from the Thalamus and Hypothalamus (Figure 7a). Since the ontology maps to the corresponding voxels of the reference space, each voxel can be related to samples that originated from the voxel’s brain region. The hierarchical nature of the ontology enables the querying of gene expression on multiple anatomical levels. For example, querying the average gene expression in the Diencephalon, the parent region of Thalamus and Hypothalamus, will aggregate over all samples of the count matrix (Figure 7b), while a query on the Thalamus or a subregion of the Thalamus (e.g. Dorsal Thalamus) will result in an aggregation over the thalamic samples (Figure 7c). We want to point out, that thalamic sample do not necessarily represent dorsal thalamic samples, hence we make the samples origin explicit in our resource’s user interface (Figure 5a, “Sample Region Annotations”).

**Fig. 7.**
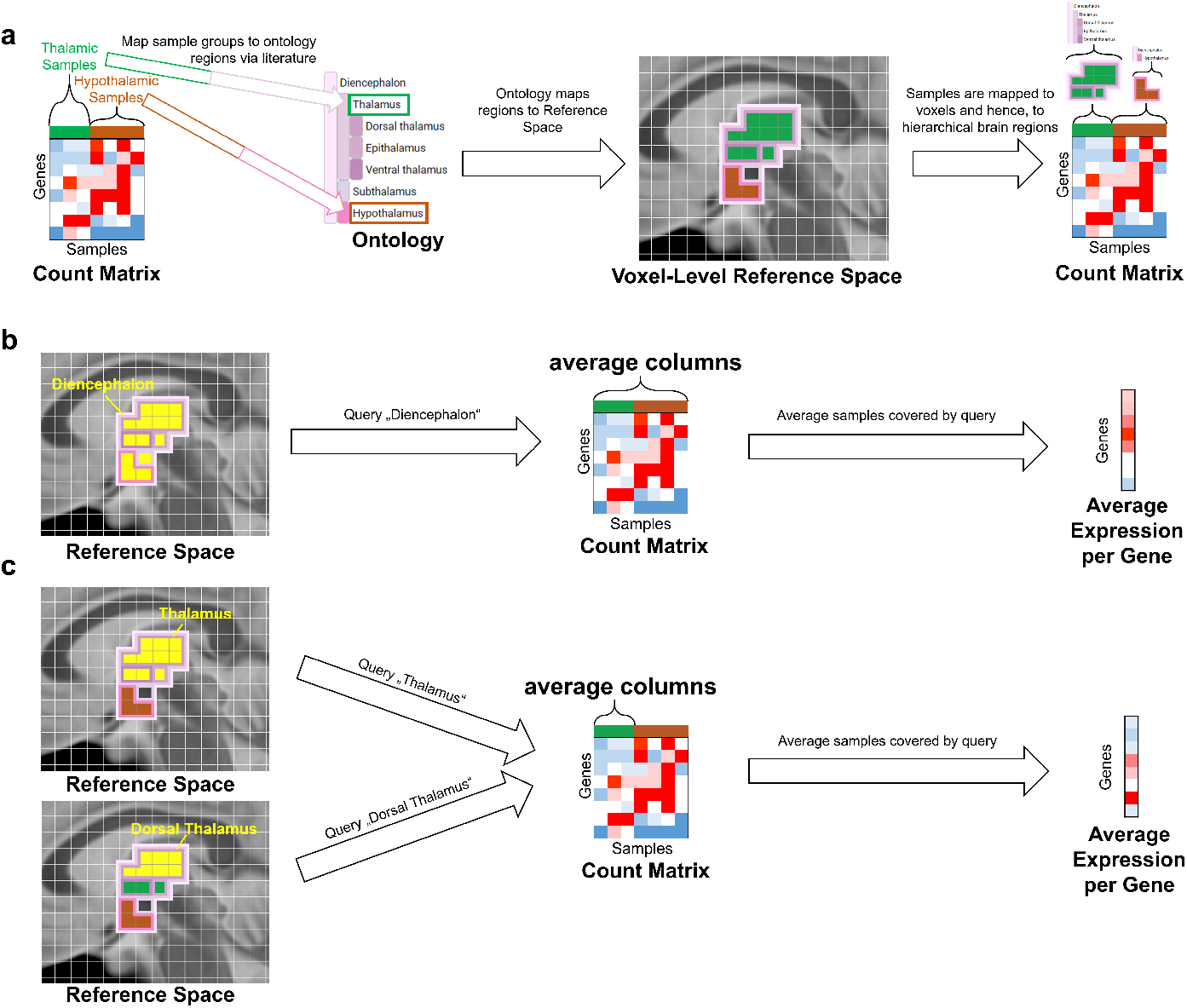
Mapping of exemplary RNA sequencing data to, and retrieving from a common reference space. a) Samples of the Thalamus (green) and Hypothalamus (brown) of an exemplary RNA sequencing count matrix are mapped manually to a brain regions in an hierarchical ontology via literature research. Since mapping of the ontology to the reference space is known, samples can be mapped to individual voxels of the reference space, and hence to every anatomical level in the ontology. b)Aggregating the average gene expression for all samples from a coarser anatomical level (Diencephalon) than the original annotations (Thalamus and Hypothalamus). c) Aggregating the average gene expression for all samples from a equal or finer anatomical level (Thalamus or Dorsal Thalamus) than the original annotations (Thalamus) leads to he same results.

We implemented four different variants of gene expression queries to cover different use cases, such as region-specificity or enrichment. Theses queries were defined on a gene expression matrix of dataset *d* as

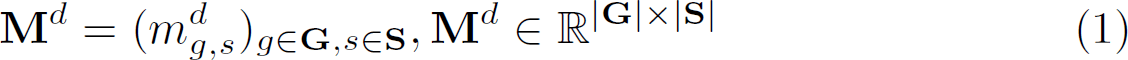

where each row represents a gene *g ∈* **G** and each column a sample (or a voxel in case of imaging data) *s ∈* **S. V** *⊆* **S** represent all samples within the *VOI*, **C** *⊆* **S** samples of a certain cell type, and **F** *⊆* **S** a samples filtered by meta data other than cell types.

- **Mean Expression Query**: Computing the mean gene expression within the *VOI* for each gene *g*

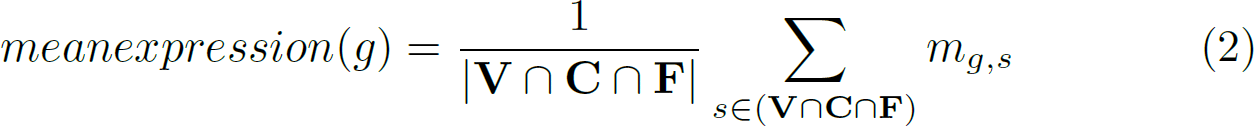
- **Region Specificity Query**: To account for regional specificity, we compute for each gene *g* the mean gene expression within the *VOI*, and normalize it to the mean expression of the rest of the brain:

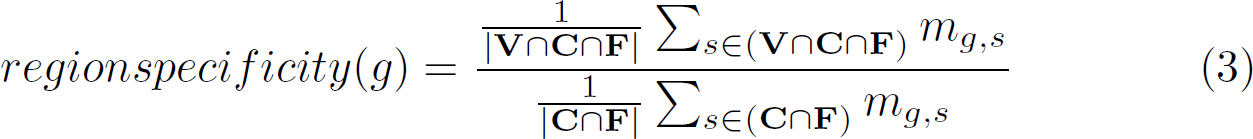
- **Cell type Specificity Query**: This query can be used to see how specific the expression of a certain cell type is. In this case, for each gene *g*, the mean gene expression within the *VOI* is computed for all samples of a certain cell type **C** *⊆* **S**, and normalized by the expression over samples of all cell types within the *VOI*:

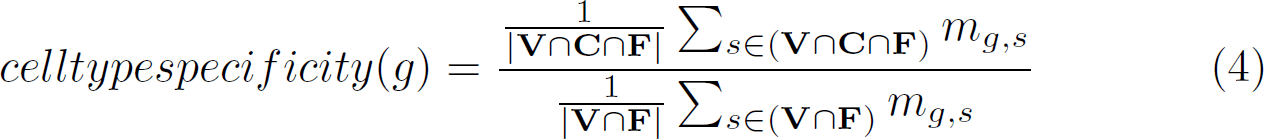
- **Enrichment Query**: This query can be used to see how specific the expression is for cell types **C** *⊆* **S** or different meta data **F** *⊆* **S**. In this case, the mean gene expression within the *VOI* is computed for all samples of the selected filter and cell type, and normalized by the expression over all samples within the *VOI*:

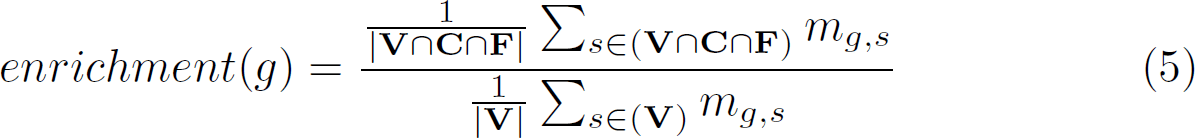

### 4.3 BrainTrawler LITE

The basis of *BrainTrawler LITE* is the dataset coverage heatmap, showing the distribution of samples/images across brain regions (columns) and datasets (rows), subdivided by meta data categories such as cell types, phenotypes etc (Figure 5a). Regions can be dynamically set via a tree-like structure showing the brain ontology (Figure 5a, left). Datasets can be subdivided by their metadata categories (e.g. split by cell types) on the right side (Figure 5a, right), so that the rows do not only represent datasets, but also subsets thereof, e.g. a row for each cell type per dataset. Hovering over the individual tiles of the heatmap reveals a summary of the data for the respective brain region and dataset (or subset of the dataset), for example sample count and the original region annotation of the data. The original region annotation is especially relevant to identify the data’s origin, and hence, the data’s potential relevance for the user.

In the first case (Figure 5b), one gene expression heatmap is generated for each of the entered genes. Here, a gene expression heatmap shows the averaged expression for all samples/images covered by each of the selected tiles in the dataset coverage heatmap. This means, that if the user selects a tile in the dataset coverage heatmap of a certain brain region, and a certain cell type of a certain dataset, each gene expression heatmap will contain the same tile, showing the averaged expression of all the tile’s covered samples/images (Figure 8a). To deal with gene lists with dozens of genes, we used a small multiples visualization [27], (Figure 5b, right) so one can visually identify patterns while maintaining an overview. Clicking on individual gene heatmaps will show a detailed view on the left hand side (Figure 5c, left), displaying the exact expression values and row/column labels. The colouring is set by individual colour scales per dataset (same colour, but the range depends on the datasets maximum value), since, as already mentioned before, values are not directly comparable across datasets.

**Fig. 8.**
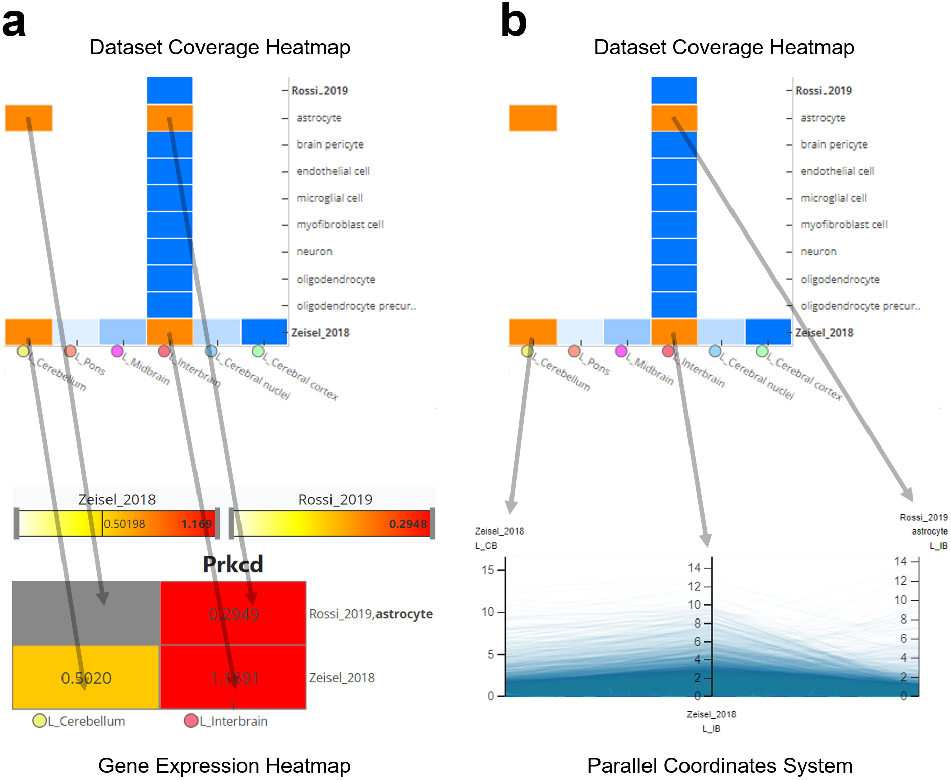
Concept of how a selection in the dataset coverage heatmap transfers to gene expression heatmaps and the parallel coordinates system. Each tile represents a subset of the resource, i.e., samples/images of a certain brain region of a certain dataset (and of a certain meta data category). Each selected tile (orange) has a direct representation as tile in the gene expression heatmap (a) and as axis in the parallel coordinates system (b). The values in (a) and (b) are the averaged expression values (e.g. CPM, TPM etc.) over all samples/images represented by the selected tile. Tiles without samples (missing data) are rendered grey in (a), or omitted for (b).

In the case of investigation on a genome level (Figure 5c), a parallel coordinates system is used analogue to the dissection of connections shown in Figure 4. Here, each line represents a gene, indicating the averaged expression along axes for each selected tile (Figure 8b). Genes can be filtered drawing brushes on an axis (Figure 5c) to find genes with specific gene expression patterns.

### 4.4 Spatial Indexing

Real-time queries on the resource’s data was achieved by spatial indexing, depending on the datatype:

- **Connectivity Data**: For real-time aggregation of connectivity data with billions of connections, we used the data structure introduced by Ganglberger et al. in 2019 [26]. Here, in principle the query speed is reached by sorting rows and columns of a connectivity matrix by their location in space along a space filling curve [88], so that rows and columns that represent connections that are close together in the 3D reference space are also close together in the matrix. This makes reading local connectivity (e.g. the connectivity of a brain region) from the hard-drive extremely efficient, since it benefits from read-ahead paging of the operating system to reach near-sequential reading speed [26].
- **Imaging Data**: For computing the mean expression of a volume of interest (*VOI*), for example a brain region, we used spatial indexing on volumetric images similar to Schulze [58]. Here, the imaging data are not stored per image, but per voxel: For each voxel in the reference space, the data of all images at the voxel’s position are stored together (i.e. on the physical harddrive). Furthermore, we order these per-voxel data along a space filling curve [88], which allows data points in close proximity in the 3D reference to be stored in close proximity as well on the storage. The expression of the voxels of a *VOI* can then be read block-wise from the hard-drive, which is more efficient than reading each image individually due to read-ahead paging of the operating system [58].
- **Sample-based Data**: For sample-based data, such as RNA sequencing and microarray gene expression data, we used a similar approach as for imaging data. Here, for each sample in our resource, we used the sample’s mapping to the reference space (Section 2.2) to get the sample’s location. Based on these locations, we order samples along a space filling curve and stored them on the hard-drive. This means, that if a certain *VOI* is queried for gene expression, all relevant samples of all datasets are stored close-together. As a consequence, they can be retrieved block-wise, benefiting from read-ahead paging of the operating system similar to the connectivity and imaging data approaches. We further optimized the queries by pre-aggregating samples with similar meta data, i.e. samples of the same dataset, cell type, age category etc. This significantly increases the query speed, since the amount of data that need to be aggregated on-the fly is reduced from thousands of individual samples to a tenth or even a hundredth of it (depending on the extend of the query).

### 4.5 Data Availability

*BrainTrawler* including the *BrainTACO* resource can be accessed after publication in a peer-reviewed journal. Code for the mapping and data generation is provided in Supplementary Data 1.

## Acknowledgments

VRVis is funded by BMK, BMDW, Styria, SFG, Tyrol and Vienna Business Agency in the scope of COMET - Competence Centers for Excellent Technologies (879730) which is managed by FFG. Wulf Haubensak was supported by the Research Institute of Molecular Pathology (IMP), Boehringer Ingelheim, the Austrian Research Promotion Agency (FFG), and a grant from the European Community’s Seventh Framework Programme (FP/2007-2013) / ERC grant agreement no. 311701. The extensions of BrainTrawler with the integration of transcriptomic data was funded by Boehringer Ingelheim, where we want to specifically thank Till Andlauer for his writing support, and Moritz von Heimendahl, Roberto Arban, Frank Gillardon, Sergio Picart-Armada, Gregiorio Alanis-Lobato, and Yasin Kaymaz for the selection and assistance with the datasets. Florian Ganglberger performed his work for the paper as Post-Doc employed at VRVis, but finalized and submitted the paper as Principal Scientist at Boehringer Ingelheim RCV GmbH & Co KG. Furthermore, we want to thank Piotr Radkowsk and Adam Filip (Ardigen) for preprocessing and Nicolas Swoboda (VRVis) for web-development support.

## 5 Supplementary information

**Supplementary Data 1:** Mapping and data generation code (related to Figure 1 and 2)

**Supplementary Data 2:** Dataset comparison code and results (related to Figure 3)

**Supplementary Table 1:** Dataset comparison figure table with additional information, including query brain regions, filters, and cell types (related to Figure 3)

**Supplementary Table 2:** Consensus hierarchy and cell types (related to Figures 6 and 6)

**Supplementary Table 3:** List of significant genes (related to Figure 6)

**Supplementary Table 4**: Species overlap (related to Figure 6)

**Supplementary Table 5:** Association summary (related to Figure 6)

**Supplementary Figure 1**: Sampled areas and connectivity analysis of AI and GI in mouse and humans (related to Figure 6)

**Supplementary Figure 2:** Overlap of genes with significantly correlated gene expression across 10 subcortical areas and AI/GI connectivity across human and mouse (related to Figure 6)

---

**Supplementary Figure 1.**
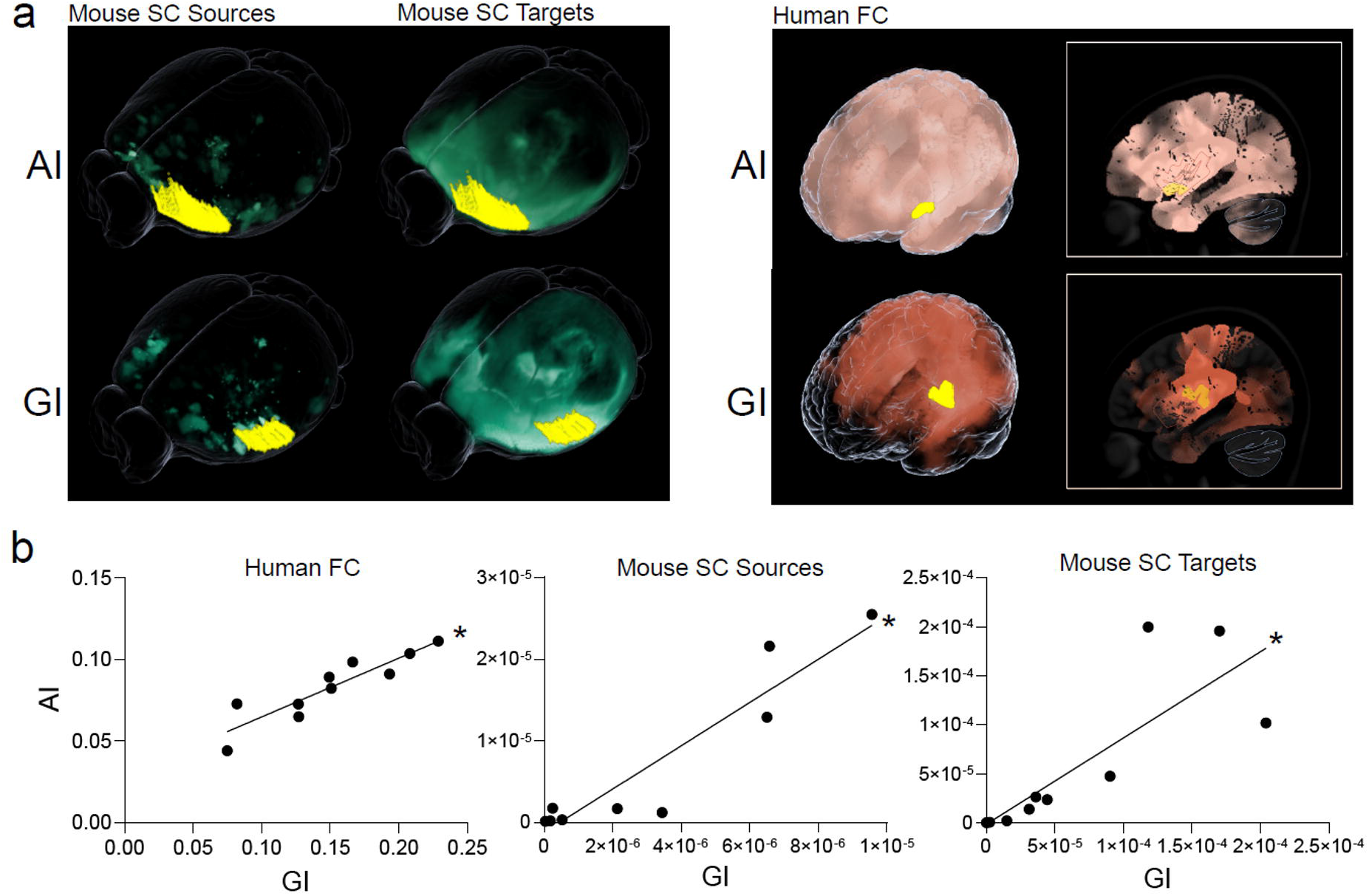
Sampled areas and connectivity analysis of Al and GI in mouse and humans. **a.** Mouse area of “L_Agranular insular area, dorsal part” and “ L_Agranular insular area, ventral part” were combined into Al, while “L_Visceral area” represents GI. Human Al and GI areas were selected by brushing sub-areas within the short and long insular gyri according to Bauernfeind et al (https://doi.org/10.1016/j.jhevol.2012.12.003). **b.** Correlations of Al and GI within Human FC (Spearman r = 0.93 p value = 0.0003) and Mouse SC Source/Targets (Spearman r = 0.92, p value = 0.0005/Spearman r = 0.94, p value = 0.0002).

**Supplementary Figure 2.**
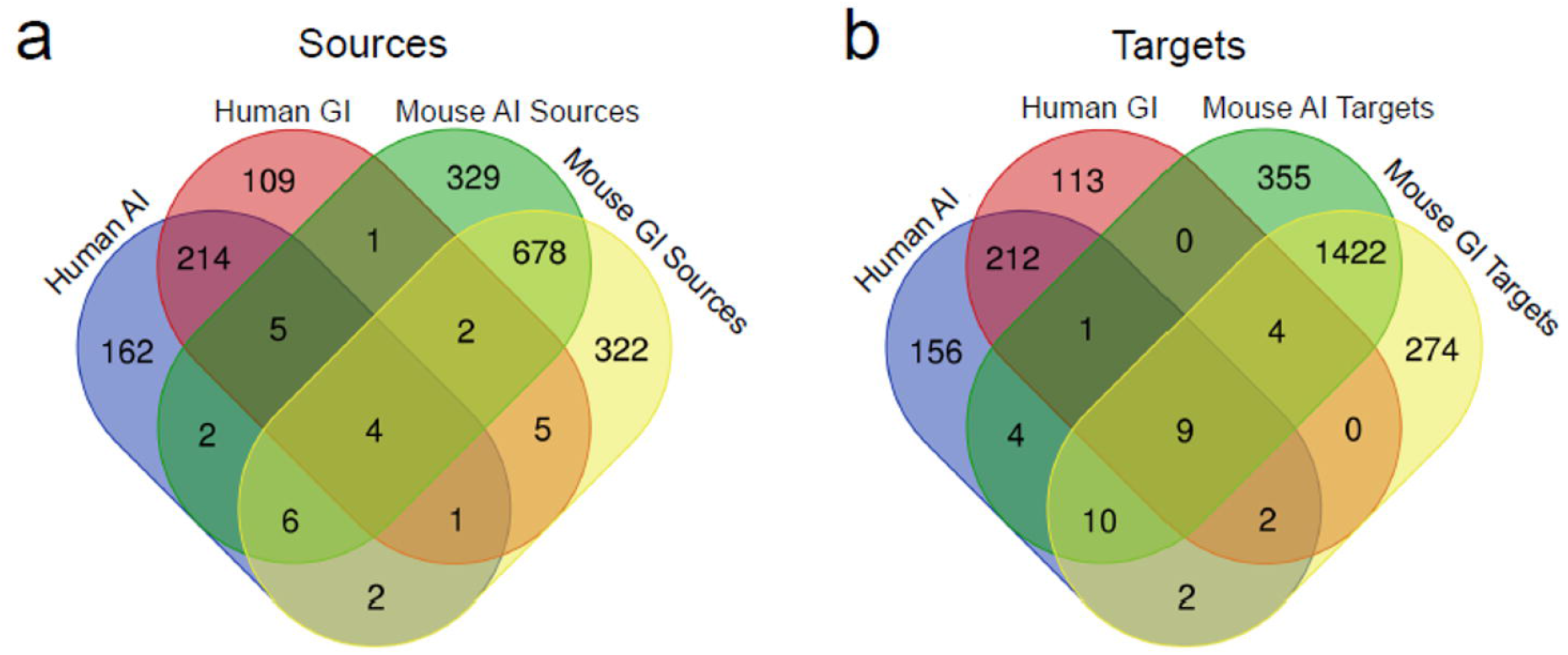
Overlap of genes with significantly correlated gene expression across 10 subcortical areas and Al/GI connectivity across human and mouse. **a.** Overlap of genes with significantly correlated gene expression across 10 subcortical areas with Human FC and Mouse SC Sources of Al and GI. **b.** Overlap of genes with significantly correlated gene expression across 10 subcortical areas with Human FC and Mouse SC Targets of Al and GI. Diagrams were generated with https://bioinformatics.psb.ugent.be/webtoolsNenn/ (See Supplementary Table 4).

