## Supplementary Data 1 for "BrainTACO: An Explorable Multi-Scale Multi-Modal Brain Transcriptomic And Connectivity Data Resource": generate_connectivity_matrix_file.R

```
# Copyright (C) 2022 VRVis.
# All rights reserved.
# Contact: VRVis Forschungs-GmbH
# 
# Redistribution and use in source and binary forms, with or without
# modification, are permitted provided that the following conditions are met:
# 1. Redistributions of source code must retain the above copyright
#    notice, this list of conditions and the following disclaimer.
# 2. Redistributions in binary form must reproduce the above copyright
#    notice, this list of conditions and the following disclaimer in the
#    documentation and/or other materials provided with the distribution.
# 3. All advertising materials mentioning features or use of this software
#    must display the following acknowledgement:
#    This product includes software developed by the VRVis Forschungs-GmbH.
# 4. Neither the name of the VRVis Forschungs-GmbH nor the
#    names of its contributors may be used to endorse or promote products
#    derived from this software without specific prior written permission.
#
# This file generates a csv connectivity matrix on voxel level (1 mm resolution) from 
# WU-Minn_​HCP_​RestingState
# run getVoxelAndVerticesToMNISpaceMapping.m in Matlab first to get the data for mapping of the voxels and vertices to the connectivity matrix but
# it's output can be found in storage/verticesAndVoxelCoordinatesFMRI.mat


library(cifti)
library(fields)
library(R.matlab)

print("Reading connectivity matrix")
fMRIConnectivityCifti<-read_cifti("HCP_S900_820_rfMRI_MSMAll_groupPCA_d4500ROW_zcorr.dconn.nii")


filename<-"HumanConnectomeProject_RestingState_voxellevel.csv"


atlasRegions<-readMat("storage/atlasRegionsHumanAHRA.mat")

index<-atlasRegions$index
indexX<-atlasRegions$indexX
indexY<-atlasRegions$indexY
indexZ<-atlasRegions$indexZ
atlasRegions<-atlasRegions$atlasRegions


#check fmri connectivity sites

fMRISites<-readMat("storage/verticesAndVoxelCoordinatesFMRI.mat")

fMRISites$verticesCoordinatesMNI[,1]<-fMRISites$verticesCoordinatesMNI[,1]+98+1;
fMRISites$verticesCoordinatesMNI[,2]<-fMRISites$verticesCoordinatesMNI[,2]+134+1;
fMRISites$verticesCoordinatesMNI[,3]<-fMRISites$verticesCoordinatesMNI[,3]+72+1;

fMRISites$verticesCoordinatesMNI<-fMRISites$verticesCoordinatesMNI[,c(2,3,1)]

fMRISites$verticesCoordinatesMNI[,1]<-dim(atlasRegions)[1]-fMRISites$verticesCoordinatesMNI[,1];
fMRISites$verticesCoordinatesMNI[,2]<-dim(atlasRegions)[2]-fMRISites$verticesCoordinatesMNI[,2];
fMRISites$verticesCoordinatesMNI[,3]<-dim(atlasRegions)[3]-fMRISites$verticesCoordinatesMNI[,3];

fMRISites$voxelCoordinatesMNI[,1]<-fMRISites$voxelCoordinatesMNI[,1]+98+1;
fMRISites$voxelCoordinatesMNI[,2]<-fMRISites$voxelCoordinatesMNI[,2]+134+1;
fMRISites$voxelCoordinatesMNI[,3]<-fMRISites$voxelCoordinatesMNI[,3]+72+1;

fMRISites$voxelCoordinatesMNI<-fMRISites$voxelCoordinatesMNI[,c(2,3,1)]

fMRISites$voxelCoordinatesMNI[,1]<-dim(atlasRegions)[1]-fMRISites$voxelCoordinatesMNI[,1];
fMRISites$voxelCoordinatesMNI[,2]<-dim(atlasRegions)[2]-fMRISites$voxelCoordinatesMNI[,2];
fMRISites$voxelCoordinatesMNI[,3]<-dim(atlasRegions)[3]-fMRISites$voxelCoordinatesMNI[,3];

  mipX<-atlasRegions[1,,]>0
  mipY<-atlasRegions[,1,]>0
  mipZ<-atlasRegions[,,1]>0
  
  
  neighbourhoodVoxels<-c(-1,0,1)
  neighbourhoodVertices<-c(-2,-1,0,1,2)
  
  for(x in 1:dim(atlasRegions)[1]){
    mipX<-mipX|atlasRegions[x,,]>0
  }
  for(actCoord in 1:dim(fMRISites$voxelCoordinatesMNI)[1]){
    mipX[fMRISites$voxelCoordinatesMNI[actCoord,2]+neighbourhoodVoxels,fMRISites$voxelCoordinatesMNI[actCoord,3]+neighbourhoodVoxels]<-2
  }
  
  for(actCoord in 1:dim(fMRISites$verticesCoordinatesMNI)[1]){
    mipX[fMRISites$verticesCoordinatesMNI[actCoord,2]+neighbourhoodVertices,fMRISites$verticesCoordinatesMNI[actCoord,3]+neighbourhoodVertices]<-2
  }
  
  image(mipX, xaxt= "n", yaxt= "n")
  
  for(y in 1:dim(atlasRegions)[2]){
    mipY<-mipY|atlasRegions[,y,]>0
  }
  for(actCoord in 1:dim(fMRISites$voxelCoordinatesMNI)[1]){
    mipY[fMRISites$voxelCoordinatesMNI[actCoord,1]+neighbourhoodVoxels,fMRISites$voxelCoordinatesMNI[actCoord,3]+neighbourhoodVoxels]<-2
  }
  
  for(actCoord in 1:dim(fMRISites$verticesCoordinatesMNI)[1]){
    mipY[fMRISites$verticesCoordinatesMNI[actCoord,1]+neighbourhoodVertices,fMRISites$verticesCoordinatesMNI[actCoord,3]+neighbourhoodVertices]<-2
  }
  
  image(mipY, xaxt= "n", yaxt= "n")
  
  for(z in 1:dim(atlasRegions)[3]){
    mipZ<-mipZ|atlasRegions[,,z]>0
  }
  for(actCoord in 1:dim(fMRISites$voxelCoordinatesMNI)[1]){
    mipZ[fMRISites$voxelCoordinatesMNI[actCoord,1]+neighbourhoodVoxels,fMRISites$voxelCoordinatesMNI[actCoord,2]+neighbourhoodVoxels]<-2
  }
  for(actCoord in 1:dim(fMRISites$verticesCoordinatesMNI)[1]){
    mipZ[fMRISites$verticesCoordinatesMNI[actCoord,1]+neighbourhoodVertices,fMRISites$verticesCoordinatesMNI[actCoord,2]+neighbourhoodVertices]<-2
  }
  
  image(mipZ, xaxt= "n", yaxt= "n")


fMRISitesIndexAtAtlas<-atlasRegions<0
for(actCoord in 1:dim(fMRISites$verticesCoordinatesMNI)[1]){
  fMRISitesIndexAtAtlas[fMRISites$verticesCoordinatesMNI[actCoord,1],fMRISites$verticesCoordinatesMNI[actCoord,2],fMRISites$verticesCoordinatesMNI[actCoord,3]]<-fMRISites$verticesIndizes[actCoord]
}
for(actCoord in 1:dim(fMRISites$voxelCoordinatesMNI)[1]){
  fMRISitesIndexAtAtlas[fMRISites$voxelCoordinatesMNI[actCoord,1],fMRISites$voxelCoordinatesMNI[actCoord,2],fMRISites$voxelCoordinatesMNI[actCoord,3]]<-fMRISites$voxelIndizes[actCoord]
}

print("Generate inflated fMRI Sites (mapping a coodinate to its closest voxel/vertex")
fMRISitesIndexAtAtlasInflated<-atlasRegions<0

pb <- txtProgressBar(min=0, max=dim(atlasRegions)[1], initial=0,style=3)
for(x in 1:dim(atlasRegions)[1]){
  setTxtProgressBar(pb, x)
  for(y in 1:dim(atlasRegions)[2]){
    for(z in 1:dim(atlasRegions)[3]){
      if(atlasRegions[x,y,z]>0){
        distancesVoxels <-rdist(fMRISites$voxelCoordinatesMNI,t(as.matrix(c(x, y, z))))
        minDistanceVoxels<-min(distancesVoxels)
        
        distancesVertices <-rdist(fMRISites$verticesCoordinatesMNI,t(as.matrix(c(x, y, z))))
        minDistanceVertices<-min(distancesVertices)
        
        if(minDistanceVoxels<=minDistanceVertices){
          fMRISitesIndexAtAtlasInflated[x,y,z]<-fMRISites$voxelIndizes[distancesVoxels==minDistanceVoxels][1]
        }else{
          fMRISitesIndexAtAtlasInflated[x,y,z]<-fMRISites$verticesIndizes[distancesVertices==minDistanceVertices][1]
        }
      }
    }
  }
}
close(pb)


  mipX<-atlasRegions[1,,]>0
  mipY<-atlasRegions[,1,]>0
  mipZ<-atlasRegions[,,1]>0
  
  for(x in 1:dim(atlasRegions)[1]){
    mipX<-mipX|atlasRegions[x,,]>0
  }
  for(x in 1:dim(atlasRegions)[1]){
    mipX[fMRISitesIndexAtAtlasInflated[x,,]>0]<-2
  }
  image(mipX, xaxt= "n", yaxt= "n")
  
  for(y in 1:dim(atlasRegions)[2]){
    mipY<-mipY|atlasRegions[,y,]>0
  }
  for(y in 1:dim(atlasRegions)[2]){
    mipY[fMRISitesIndexAtAtlasInflated[,y,]>0]<-2
  }
  image(mipY, xaxt= "n", yaxt= "n")
  
  for(z in 1:dim(atlasRegions)[3]){
    mipZ<-mipZ|atlasRegions[,,z]>0
  }
  for(z in 1:dim(atlasRegions)[3]){
    mipZ[fMRISitesIndexAtAtlasInflated[,,z]>0]<-2
  }
  image(mipZ, xaxt= "n", yaxt= "n")


print("Generate hilbert order")
hilbert3d<-function(n){
  if(n <= 0){
    x<-0;
    y<-0;
    z<-0;
  }else{
    lookup<-hilbert3d(n-1);
    x = 0.5*c(0.5+lookup$z , 0.5+lookup$y,-0.5+lookup$y,-0.5-lookup$x,-0.5-lookup$x,-0.5-lookup$y, 0.5-lookup$y, 0.5+lookup$z);
    y = 0.5*c(0.5+lookup$x , 0.5+lookup$z,0.5+lookup$z,0.5+lookup$y,-0.5+lookup$y,-0.5-lookup$z, -0.5-lookup$z, -0.5-lookup$x);
    z = 0.5*c(0.5+lookup$y , -0.5+lookup$x,-0.5+lookup$x,0.5-lookup$z,0.5-lookup$z,-0.5+lookup$x, -0.5+lookup$x, 0.5-lookup$y);
  }
  
  return(data.frame(x=x,y=y,z=z))
}

orderN<-1
lookup<-data.frame(x=0,y=0,z=0)

while((max(indexX[atlasRegions>0])-min(indexX[atlasRegions>0]))>=(2^orderN) || (max(indexY[atlasRegions>0])-min(indexY[atlasRegions>0]))>=(2^orderN) || (max(indexZ[atlasRegions>0])-min(indexZ[atlasRegions>0]))>=(2^orderN)){
  orderN<-orderN+1
}

lookup<-ceiling((hilbert3d(orderN)+0.5)*(2^orderN))
lookup$x<-lookup$x-1+min(indexX[atlasRegions>0])
lookup$y<-lookup$y-1+min(indexY[atlasRegions>0])
lookup$z<-lookup$z-1+min(indexZ[atlasRegions>0])

lookup<-lookup[lookup$x<=max(indexX) & lookup$y<=max(indexY) & lookup$z<=max(indexZ),]


indizesInHilbertOrder<-rep(0,dim(lookup)[1])
hilbertsIndex<-1:dim(lookup)[1]
for(actHilbertIndex in hilbertsIndex){
  indizesInHilbertOrder[actHilbertIndex]<-index[lookup$x[actHilbertIndex],lookup$y[actHilbertIndex],lookup$z[actHilbertIndex]]
  if(actHilbertIndex%%10000 == 0){
    print(paste0(actHilbertIndex/length(hilbertsIndex)*100,"%"))
  }
}
print(paste0("100%"))

hilbertfMRISitesIndizes<-fMRISitesIndexAtAtlas
hilbertfMRISitesIndizes<-hilbertfMRISitesIndizes[indizesInHilbertOrder]
hilbertfMRISitesIndizes<-hilbertfMRISitesIndizes[hilbertfMRISitesIndizes>0]

matrixIndex<-1:length(hilbertfMRISitesIndizes)
matrixNames<-rep(NA,length(hilbertfMRISitesIndizes))
matrixNamesOrg<-hilbertfMRISitesIndizes


print("Mapping connectivity voxels/vertices to connectivity matrix rows/column names")
pb <- txtProgressBar(min=0, max=100, initial=0,style=3)
for(actNameIndex in 1:length(matrixNames)){
  setTxtProgressBar(pb, actNameIndex/(length(matrixNames))*100)
  indexOfActName<-fMRISitesIndexAtAtlasInflated==hilbertfMRISitesIndizes[actNameIndex]
  if(sum(indexOfActName)>0){
    matrixNames[actNameIndex]<-paste0(index[cbind(indexX[indexOfActName],indexY[indexOfActName],indexZ[indexOfActName])],collapse="_")
  }
}
close(pb)
sum(is.na(matrixNames))

options(scipen=20)

indizesToAdd<-matrixNamesOrg[!is.na(matrixNames)]


  print("Creating connectivity matrix")
  write.table(format(t(as.matrix(c(0,matrixNames[!is.na(matrixNames)]))), scientific=FALSE), file=filename, append=FALSE,row.names=F, col.names=F,  sep=",", quote=F)
  
  act<-1
  pb <- txtProgressBar(min=0, max=100, initial=0,style=3)
  for(actIndex in indizesToAdd){
    a<-(t(as.matrix(fMRIConnectivityCifti$data[actIndex,indizesToAdd])))
    a<-round(a,digits=3)
    rownames(a)<-format(matrixNames[actIndex==matrixNamesOrg],scientific=FALSE)
    write.table(formatC(a,  format="fg"), file=filename, append=TRUE,row.names=T, col.names=F,  sep=",", quote=F)
    
    act=act+1;
    setTxtProgressBar(pb, act/(length(indizesToAdd))*100)
  }
  
  close(pb)
```
