## Supplementary Data 1 for "BrainTACO: An Explorable Multi-Scale Multi-Modal Brain Transcriptomic And Connectivity Data Resource": generateReferenceSpace.R

```
# Copyright (C) 2022 VRVis.
# All rights reserved.
# Contact: VRVis Forschungs-GmbH
# 
# Redistribution and use in source and binary forms, with or without
# modification, are permitted provided that the following conditions are met:
# 1. Redistributions of source code must retain the above copyright
#    notice, this list of conditions and the following disclaimer.
# 2. Redistributions in binary form must reproduce the above copyright
#    notice, this list of conditions and the following disclaimer in the
#    documentation and/or other materials provided with the distribution.
# 3. All advertising materials mentioning features or use of this software
#    must display the following acknowledgement:
#    This product includes software developed by the VRVis Forschungs-GmbH.
# 4. Neither the name of the VRVis Forschungs-GmbH nor the
#    names of its contributors may be used to endorse or promote products
#    derived from this software without specific prior written permission.
#
# This file generates a voxel level reference space with region annotations on a 1 mm 152 MNI space


library(RNifti)
library(fields)
library(R.matlab)
library(icesTAF) #Needed to create amira files
library(R.utils)

dir.create("storage") 
download.file("http://download.alleninstitute.org/informatics-archive/allen_human_reference_atlas_3d_2020/version_1/annotation.nii.gz", "annotation.nii.gz")
gunzip("annotation.nii.gz","annotation.nii")
file.rename("annotation.nii","storage\\annotation.nii")
atlasRegions_org<-readNifti("storage\\annotation.nii")
atlasRegions_header<-niftiHeader("storage\\annotation.nii")

for(x in 1:floor(dim(atlasRegions_org)[1]/2)){
  atlasRegions_org[dim(atlasRegions_org)[1]-(x-1),,]<-atlasRegions_org[x,,]
}
image(atlasRegions_org[330,,]>0, xaxt= "n", yaxt= "n")


voxelsize<-1000

#downsample the data to 1 voxel per mm
atlasRegions<-array(0, dim(atlasRegions_org)/2); 
Mode <- function(x) {
  ux <- unique(x[!is.na(x) & x>0])
  if(length(ux)==0){
    return(0)
  }
  ux[which.max(tabulate(match(x[!is.na(x) & x>0], ux)))]
}

for(x in 1:(dim(atlasRegions)[1])){
  for(y in 1:(dim(atlasRegions)[2])){
    for(z in 1:(dim(atlasRegions)[3])){
      atlasRegions[x,y,z]<-Mode(as.vector(atlasRegions_org[((2*x-1):(2*x)),((2*y-1):(2*y)),((2*z-1):(2*z))]))
    }
  }
}

#mirror data (data is only on one side)
for(x in 1:floor(dim(atlasRegions)[1]/2)){
  atlasRegions[dim(atlasRegions)[1]-(x-1),,]<-atlasRegions[x,,]
}

atlasRegionsPerm <- aperm(atlasRegions, c(2,3,1))
atlasRegionsPerm_flipped<-atlasRegionsPerm
for(x in 1:dim(atlasRegionsPerm)[1]){
  atlasRegionsPerm_flipped[dim(atlasRegionsPerm)[1]-(x-1),,]<-atlasRegionsPerm[x,,]
}
atlasRegionsPerm<-atlasRegionsPerm_flipped

atlasRegionsPerm_flipped<-atlasRegionsPerm
for(y in 1:dim(atlasRegionsPerm)[2]){
  atlasRegionsPerm_flipped[,dim(atlasRegionsPerm)[2]-(y-1),]<-atlasRegionsPerm[,y,]
}
atlasRegionsPerm<-atlasRegionsPerm_flipped

atlasRegionsPerm_flipped<-atlasRegionsPerm
for(z in 1:dim(atlasRegionsPerm)[3]){
  atlasRegionsPerm_flipped[,,dim(atlasRegionsPerm)[3]-(z-1)]<-atlasRegionsPerm[,,z]
}
atlasRegionsPerm<-atlasRegionsPerm_flipped

atlasRegions<-atlasRegionsPerm


image(atlasRegions[,100,]>0, xaxt= "n", yaxt= "n")

image(atlasRegions[,,100]>0, xaxt= "n", yaxt= "n")

image(atlasRegions[100,,]>0, xaxt= "n", yaxt= "n")

length(unique(as.vector(atlasRegions_org)))
length(unique(as.vector(atlasRegions)))


index<-atlasRegions
index[1:length(as.vector(atlasRegions))]<- 1:length(as.vector(atlasRegions))
atlasRegionsBiggerZero<-atlasRegions[atlasRegions>0]

sum(atlasRegions>0)

indexX=rep(0,length(atlasRegions))
indexY=rep(0,length(atlasRegions))
indexZ=rep(0,length(atlasRegions))

for(x in 1:(dim(index)[1])){
  for(y in 1:(dim(index)[2])){
    for(z in 1:(dim(index)[3])){
      indexX[index[x,y,z]]=x
      indexY[index[x,y,z]]=y
      indexZ[index[x,y,z]]=z
    }
  }
}

writeMat("atlasRegionsHumanAHRA.mat",atlasRegions=atlasRegions,index=index,indexX=indexX,indexY=indexY,indexZ=indexZ)
file.rename("atlasRegionsHumanAHRA.mat","storage\\atlasRegionsHumanAHRA.mat")
```
