## Supplementary Data 1 for "BrainTACO: An Explorable Multi-Scale Multi-Modal Brain Transcriptomic And Connectivity Data Resource": getVoxelAndVerticesToMNISpaceMapping.m

```
% Copyright (C) 2022 VRVis.
% All rights reserved.
% Contact: VRVis Forschungs-GmbH
% 
% Redistribution and use in source and binary forms, with or without
% modification, are permitted provided that the following conditions are met:
% 1. Redistributions of source code must retain the above copyright
%    notice, this list of conditions and the following disclaimer.
% 2. Redistributions in binary form must reproduce the above copyright
%    notice, this list of conditions and the following disclaimer in the
%    documentation and/or other materials provided with the distribution.
% 3. All advertising materials mentioning features or use of this software
%    must display the following acknowledgement:
%    This product includes software developed by the VRVis Forschungs-GmbH.
% 4. Neither the name of the VRVis Forschungs-GmbH nor the
%    names of its contributors may be used to endorse or promote products
%    derived from this software without specific prior written permission.
%
% This function processes the grayordinate functional resting connectivity data from the
% WU-Minn Human Connectome Project and converts it to voxel level ICBM 152 MNI space. 
% Note that you need to apply for access and download the grayordinate based data HCP_S900_820_rfMRI_MSMAll_groupPCA_d4500ROW_zcorr.dconn.nii from 
% https://www.humanconnectome.org/
%
% For the connectome workbench, run 
% export LD_LIBRARY_PATH=/home/users/ganglberger/workbench-linux64-v1.1.1/workbench/libs_linux64
% before starting matlab


conFilename = 'HCP_S900_820_rfMRI_MSMAll_groupPCA_d4500ROW_zcorr.dconn.nii';
cortexLeftFile = 'S900.L.white_MSMAll.32k_fs_LR.surf.gii';
cortexRightFile = 'S900.R.white_MSMAll.32k_fs_LR.surf.gii';


[status,cmdout] = system(['/home/users/ganglberger/workbench-linux64-v1.1.1/workbench/exe_linux64/wb_command' ' -file-information ' conFilename]);
cmdout

[status,cmdout] = system(['/home/users/ganglberger/workbench-linux64-v1.1.1/workbench/exe_linux64/wb_command' ' -file-information ' cortexRightFile]);
cmdout

%transformation to MNI space
connectivityPosToMNItransform = [-2,0,0,90;0,2,0,-126;0,0,2,-72;0,0,0,1];


%get the voxel coordinates of the subcortical voxels
[status,cmdout] = system(['/home/users/ganglberger/workbench-linux64-v1.1.1/workbench/exe_linux64/wb_command' ' -cifti-export-dense-mapping ' conFilename ' ROW -volume-all voxelIndizes.txt']);
[status,cmdout] = system(['/home/users/ganglberger/workbench-linux64-v1.1.1/workbench/exe_linux64/wb_command' ' -cifti-export-dense-mapping ' conFilename ' ROW -surface CORTEX_LEFT verticesIndizesLeft.txt']);
[status,cmdout] = system(['/home/users/ganglberger/workbench-linux64-v1.1.1/workbench/exe_linux64/wb_command' ' -cifti-export-dense-mapping ' conFilename ' ROW -surface CORTEX_RIGHT verticesIndizesRight.txt']);

%get surface vertices coordinates 
[status,cmdout] = system(['/home/users/ganglberger/workbench-linux64-v1.1.1/workbench/exe_linux64/wb_command' ' -surface-coordinates-to-metric ' cortexLeftFile ' coordinatesVertices_left.gii']);
[status,cmdout] = system(['/home/users/ganglberger/workbench-linux64-v1.1.1/workbench/exe_linux64/wb_command' ' -surface-coordinates-to-metric ' cortexRightFile ' coordinatesVertices_right.gii']);


 fid = fopen('voxelIndizes.txt');
 indizesFile = textscan(fid,'%f%f%f%f','delimiter',' ');
 fclose(fid);
 
 %voxel coordiantes to MNI space
 voxelCoordinatesMNI = (connectivityPosToMNItransform * [[indizesFile{2},indizesFile{3},indizesFile{4}]'; ones(1, size(indizesFile{1},1))])';
 voxelCoordinatesMNI=voxelCoordinatesMNI(1:end,1:3);
 voxelIndizes=indizesFile{1}+1;


 %vertices coordinates to MNI space
 verticesIndizes = [];
 verticesCoordinatesMNI = [];
 
 cFile = gifti('coordinatesVertices_left.gii');
 
 fid = fopen('verticesIndizesLeft.txt');
 indizesFile = textscan(fid,'%f%f','delimiter',' ');
 fclose(fid);
 verticesIndizes = [verticesIndizes;indizesFile{1}+1];
 verticesCoordinatesMNI = [verticesCoordinatesMNI;cFile.cdata(indizesFile{2}+1,:)];

 cFile = gifti('coordinatesVertices_right.gii');
 
 fid = fopen('verticesIndizesRight.txt');
 indizesFile = textscan(fid,'%f%f','delimiter',' ');
 fclose(fid);
 verticesIndizes = [verticesIndizes;indizesFile{1}+1];
 verticesCoordinatesMNI = [verticesCoordinatesMNI;cFile.cdata(indizesFile{2}+1,:)];
 
 %save this data so you can cross check it in R
  save('verticesAndVoxelCoordinatesFMRI.mat','verticesCoordinatesMNI');
  save('verticesAndVoxelCoordinatesFMRI.mat','voxelCoordinatesMNI','-append');
  save('verticesAndVoxelCoordinatesFMRI.mat','voxelIndizes','-append');
  save('verticesAndVoxelCoordinatesFMRI.mat','verticesIndizes','-append');
```
