## Supplementary Data 1 for "BrainTACO: An Explorable Multi-Scale Multi-Modal Brain Transcriptomic And Connectivity Data Resource": generate_connectivity_matrix_file.R

```
# Copyright (C) 2022 VRVis.
# All rights reserved.
# Contact: VRVis Forschungs-GmbH
# 
# Redistribution and use in source and binary forms, with or without
# modification, are permitted provided that the following conditions are met:
# 1. Redistributions of source code must retain the above copyright
#    notice, this list of conditions and the following disclaimer.
# 2. Redistributions in binary form must reproduce the above copyright
#    notice, this list of conditions and the following disclaimer in the
#    documentation and/or other materials provided with the distribution.
# 3. All advertising materials mentioning features or use of this software
#    must display the following acknowledgement:
#    This product includes software developed by the VRVis Forschungs-GmbH.
# 4. Neither the name of the VRVis Forschungs-GmbH nor the
#    names of its contributors may be used to endorse or promote products
#    derived from this software without specific prior written permission.
#
# This file generates a csv connectivity matrix on voxel level (100 micron resolution) from 
# Allen Mouse brain axonal projections
# run get_connectivity_data.m in Matlab first to get the data for generating the connectivity matrix

library(R.matlab)
library(fields)
library(rjson)
library(Matrix)

generateEdgeListFile<-FALSE
generateNear<-FALSE

index<-c()

atlasRegions<-readMat("storage//atlasRegions.mat")
index<-atlasRegions$index
atlasRegions<-as.vector(atlasRegions$atlasRegions)
voxelsize<-100

atlasRegionsBiggerZero<-atlasRegions[atlasRegions>0]

sum(atlasRegions>0)

indexX=c()
indexY=c()
indexZ=c()

for(x in 1:(dim(index)[1])){
  for(y in 1:(dim(index)[2])){
    for(z in 1:(dim(index)[3])){
      indexX[index[x,y,z]]=x
      indexY[index[x,y,z]]=y
      indexZ[index[x,y,z]]=z
    }
  }
}

mY<-max(indexY)
mZ<-max(indexZ)
filename<-""

conData<-readMat("storage//injectedIndizes.mat")
filename<-"F:\\rawconnectivitydata\\AMBAConMouseBrainHilbert.csv"


injectedIDs<-unique(as.vector(conData$injectedIDs))
injectedIDsIndex<-c()
injectedIDsIndex[injectedIDs]<-1:length(injectedIDs)

injectedIndizesToID<-((conData$injectedIndizesToID))
injectedIndizes<-unique(injectedIndizesToID[,1])

injectedIndizesIndex<-c()
injectedIndizesIndex[injectedIndizes]<-1:length(injectedIndizes)
indexMatrix<-as.vector(conData$indexMatrix)
remove(conData)

length(unique(injectedIDs))
length(injectedIndizes)

hilbert3d<-function(n){
  if(n <= 0){
    x<-0;
    y<-0;
    z<-0;
  }else{
    lookup<-hilbert3d(n-1);
    x = 0.5*c(0.5+lookup$z , 0.5+lookup$y,-0.5+lookup$y,-0.5-lookup$x,-0.5-lookup$x,-0.5-lookup$y, 0.5-lookup$y, 0.5+lookup$z);
    y = 0.5*c(0.5+lookup$x , 0.5+lookup$z,0.5+lookup$z,0.5+lookup$y,-0.5+lookup$y,-0.5-lookup$z, -0.5-lookup$z, -0.5-lookup$x);
    z = 0.5*c(0.5+lookup$y , -0.5+lookup$x,-0.5+lookup$x,0.5-lookup$z,0.5-lookup$z,-0.5+lookup$x, -0.5+lookup$x, 0.5-lookup$y);
  }
  
  return(data.frame(x=x,y=y,z=z))
}

orderN<-1
lookup<-data.frame(x=0,y=0,z=0)

while((max(indexX[atlasRegions>0])-min(indexX[atlasRegions>0]))>=(2^orderN) && (max(indexY[atlasRegions>0])-min(indexY[atlasRegions>0]))>=(2^orderN) && (max(indexZ[atlasRegions>0])-min(indexZ[atlasRegions>0]))>=(2^orderN)){

  orderN<-orderN+1
}

orderN<-orderN+1
lookup<-ceiling((hilbert3d(orderN)+0.5)*(2^orderN))
lookup$x<-lookup$x-1+min(indexX[atlasRegions>0])
lookup$y<-lookup$y-1+min(indexY[atlasRegions>0])
lookup$z<-lookup$z-1+min(indexZ[atlasRegions>0])

lookup<-lookup[lookup$x<=max(indexX) & lookup$y<=max(indexY) & lookup$z<=max(indexZ),]


indizesInHilbertOrder<-rep(0,dim(lookup)[1])
hilbertsIndex<-1:dim(lookup)[1]
for(actHilbertIndex in hilbertsIndex){
  indizesInHilbertOrder[actHilbertIndex]<-index[lookup$x[actHilbertIndex],lookup$y[actHilbertIndex],lookup$z[actHilbertIndex]]
  if(actHilbertIndex%%10000 == 0){
    print(paste0(actHilbertIndex/length(hilbertsIndex)*100,"%"))
  }
}
print(paste0("100%"))


injectedIndizesInHilbertOrder<-indizesInHilbertOrder[is.element(indizesInHilbertOrder,injectedIndizes)]


hilbertatlasIndizesWithinBrain<-rep(0,length(atlasRegions)) 
hilbertatlasIndizesWithinBrain[atlasRegions>0]<-1:sum(atlasRegions>0)
hilbertatlasIndizesWithinBrain<-hilbertatlasIndizesWithinBrain[indizesInHilbertOrder]
hilbertatlasIndizesWithinBrain<-hilbertatlasIndizesWithinBrain[hilbertatlasIndizesWithinBrain>0]


pb <- txtProgressBar(min=0, max=100, initial=0,style=3)
act=1;


matrixIndex<-1:sum(atlasRegions>0)
matrixNamesOrg<-(1:length(atlasRegions))[atlasRegions>0]
matrixNames<-(1:length(atlasRegions))[atlasRegions>0]

write.table(format(t(as.matrix(c(0,matrixNames[hilbertatlasIndizesWithinBrain]))), scientific=FALSE), file=filename, append=FALSE,row.names=F, col.names=F,  sep=",", quote=F)


indizesToAdd<-injectedIndizesInHilbertOrder[atlasRegions[injectedIndizesInHilbertOrder]>0]

for(actIndex in indizesToAdd){
  
    idSofIndex<-unique(injectedIndizesToID[injectedIndizesToID[,1]==actIndex,2])
    
    conRow<-c()
    
    #load connectivity from hd
    actRow<-0
    for(i in injectedIDsIndex[idSofIndex]){
      conInd<-readMat(paste0("storage//id_connectivity/",i,".mat"),sep="")
      
      conRowHelp<-(as.vector(conInd$con))[atlasRegions>0]
      conRowHelp[conRowHelp<0]<-0
      
      conRowHelp[is.nan(conRowHelp)]<-0
      conRowHelp[conRowHelp==Inf]<-0
      conRowHelp[conRowHelp<10^-4.5]<-0
      
      if(actRow==0){
        conRow<-conRowHelp
      }else{
        conRow<-pmax(conRow,conRowHelp)
      }
      
      actRow=actRow+1;
   
    }
  
   rowData<-(t(as.matrix(conRow[hilbertatlasIndizesWithinBrain])))
   rownames(rowData)<-format(matrixNames[actIndex==matrixNamesOrg],scientific=FALSE)
    
   write.table(formatC(rowData,  format="fg", digits=5), file=filename, append=TRUE,row.names=T, col.names=F,  sep=",", quote=F)
  
   act=act+1;
   setTxtProgressBar(pb, act/(length(indizesToAdd))*100)
  
}

close(pb)
```
