## Supplementary Data 1 for "BrainTACO: An Explorable Multi-Scale Multi-Modal Brain Transcriptomic And Connectivity Data Resource": get_connectivity_data.m

```
% Copyright (C) 2022 VRVis.
% All rights reserved.
% Contact: VRVis Forschungs-GmbH
% 
% Redistribution and use in source and binary forms, with or without
% modification, are permitted provided that the following conditions are met:
% 1. Redistributions of source code must retain the above copyright
%    notice, this list of conditions and the following disclaimer.
% 2. Redistributions in binary form must reproduce the above copyright
%    notice, this list of conditions and the following disclaimer in the
%    documentation and/or other materials provided with the distribution.
% 3. All advertising materials mentioning features or use of this software
%    must display the following acknowledgement:
%    This product includes software developed by the VRVis Forschungs-GmbH.
% 4. Neither the name of the VRVis Forschungs-GmbH nor the
%    names of its contributors may be used to endorse or promote products
%    derived from this software without specific prior written permission.
%
% This script downloads the injection based connectivity data from the
% Allen Brain Atlas API. Every downloaded injection-file represents the
% connectivity to the target regions (grid points) of an injection. The
% source grid points are stored into storage/injectedIndizes.mat and
% correspond to the filenames of the injection files 
% (storage/id_connectivity) named by a continuous id.
% The injections are mirrored to both hemispheres. The connectivity of
% every injection image is normalized by its injection volume.
%

    %download the annotation of the Allen Mouse Common Coordinate Framework
    %in this case, only because we need the dimension-size of the space
    urlwrite('http://download.alleninstitute.org/informatics-archive/current-release/mouse_ccf/annotation/ccf_2015/annotation_100.nrrd','storage/annotation_100.nrrd');
    [newANOGD, metaDMASK] = nrrdread('storage/annotation_100.nrrd');
    ANOGD=permute(newANOGD,[2 1 3]);

    index=reshape(1:(size(ANOGD,1)*size(ANOGD,2)*size(ANOGD,3)),size(ANOGD,1),size(ANOGD,2),size(ANOGD,3));

    atlasRegions = ANOGD;
    save('storage//atlasRegions.mat','atlasRegions');
    save('storage//atlasRegions.mat','index','-append');


    indexX=1:(size(ANOGD,1)*size(ANOGD,2)*size(ANOGD,3));
    indexY=1:(size(ANOGD,1)*size(ANOGD,2)*size(ANOGD,3));
    indexZ=1:(size(ANOGD,1)*size(ANOGD,2)*size(ANOGD,3));

    for x= 1:size(ANOGD,1)
        for y= 1:size(ANOGD,2)
            for z= 1:size(ANOGD,3)
                indexX(index(x,y,z))=x;
                indexY(index(x,y,z))=y;
                indexZ(index(x,y,z))=z;
            end
        end
    end

    injectedIDs = [];
    injectedIndizesToID= [];

    injCount=1;

    mkdir('storage//id_connectivity');
    mkdir('storage//injection_raw_data');
    
    disp('Get injection coordinates from API')
    urlwrite('http://api.brain-map.org/api/v2/data/query.xml?criteria=service::mouse_connectivity_injection_structure[primary_structure_only$eqtrue]', 'storage/injections.xml');
    root = xmlread('storage/injections.xml');
    
    totalAmount = str2num(root.item(0).getAttribute('num_rows'));

    %traverse injections from storage/injections.xml
    for actSectionDataSet=0:root.item(0).item(0).getChildNodes.getLength-1;
        selectionDataSet = root.item(0).item(0).getChildNodes.item(actSectionDataSet);
        if strcmp(selectionDataSet,'[object: null]')
            id = -1;
            injectionX=-1;
            injectionY=-1;
            injectionZ=-1;

            [ attribute, childnodes ] = goToChildNode( selectionDataSet,'id',0 );
            id = str2num(attribute);

            [ attribute, injectionCoordinates ] = goToChildNode( selectionDataSet,'injection-coordinates',0 );
            [ attribute, childnodes ] = goToChildNode( injectionCoordinates,'injection-coordinate',0 );
            injectionX = str2num(attribute)/100+0;
            [ attribute, childnodes ] = goToChildNode( injectionCoordinates,'injection-coordinate',1 );
            injectionY = str2num(attribute)/100+0;
            [ attribute, childnodes ] = goToChildNode( injectionCoordinates,'injection-coordinate',2 );
            injectionZ = str2num(attribute)/100+0;


          
            injectionX=round(injectionX);
            injectionY=round(injectionY);
            injectionZ=round(injectionZ);

            if id>-1
                found=false;
                trying=0;
                while found==false
                    try
                        if((not(exist(sprintf('storage\\injection_raw_data\\%d.nrrd',id), 'file')) ...
                            || not(exist(sprintf('storage\\injection_raw_data\\%d_inj.nrrd',id), 'file')) ...
                            || not(exist(sprintf('storage\\injection_raw_data\\%d_inj_f.nrrd',id), 'file')) ...
                            || not(exist(sprintf('storage\\injection_raw_data\\%d_mask.nrrd',id), 'file'))) || trying>0)
                        
                            disp(sprintf('   Download data for injection %d of %d...',(injCount+1)/2,totalAmount))
                            urlwrite(sprintf('http://api.brain-map.org/grid_data/download_file/%d?image=projection_density&resolution=100',id), sprintf('storage\\injection_raw_data\\%d.nrrd',id));
                            urlwrite(sprintf('http://api.brain-map.org/grid_data/download_file/%d?image=injection_density&resolution=100',id), sprintf('storage\\injection_raw_data\\%d_inj.nrrd',id));
                            urlwrite(sprintf('http://api.brain-map.org/grid_data/download_file/%d?image=injection_fraction&resolution=100',id), sprintf('storage\\injection_raw_data\\%d_inj_f.nrrd',id));
                            urlwrite(sprintf('http://api.brain-map.org/grid_data/download_file/%d?image=data_mask&resolution=100',id), sprintf('storage\\injection_raw_data\\%d_mask.nrrd',id));
                        else
                            disp(sprintf('   Load local data for injection %d of %d...',(injCount+1)/2,totalAmount))
                        end

                        [injectionImg, metaDMASK] = nrrdread(sprintf('storage\\injection_raw_data\\%d.nrrd',id));
                        [DMASK, metaDMASK] = nrrdread(sprintf('storage\\injection_raw_data\\%d_mask.nrrd',id));

                        injectionImg(not(DMASK==1))=0;
                        injectionImg(injectionImg<0)=0;
                        injectionImg=permute(injectionImg,[2 1 3]);

                        [injectionSite, metaDMASK] = nrrdread(sprintf('storage\\injection_raw_data\\%d_inj_f.nrrd',id));
                        injectionSite(not(DMASK==1))=0;
                        [injectionVolume, metaDMASK] = nrrdread(sprintf('storage\\injection_raw_data\\%d_inj.nrrd',id));
                        injectionVolume(not(DMASK==1))=0;
                        injectionVolume=sum(injectionVolume(:));

                        injectionSite=permute(injectionSite,[2 1 3]);

                        injectionImg(injectionSite>=1)=0;

                        found=true;
                       
                    catch err
                        pause(1)
                        trying=trying+1;
                        disp(err.message)
                        disp(sprintf('Problem after download, retrying %d',trying))
                        found=false;
                    end
                end


                con=reshape(injectionImg/injectionVolume,size(ANOGD,1)*size(ANOGD,2)*size(ANOGD,3),1);
                save(sprintf('storage\\id_connectivity\\%d',injCount),'con');
              

                %add the indizes of the source grid points to injectedIndizesToID
                injectedIDs=[injectedIDs,injCount];
                injectedIndizesToID(size(injectedIndizesToID,1)+1,:)=[index(injectionX,injectionY,injectionZ),injCount];
                for i = index(injectionSite>=1.0)'
                    injectedIndizesToID(size(injectedIndizesToID,1)+1,:)=[index(indexX(i),indexY(i),indexZ(i)),injCount];
                end

                injCount=injCount+1;

                %Mirror Z-Axis, so that the injection will be mirrored to
                %the other hemisphere
                injectionImg = injectionImg(:,:,end:-1:1);
                injectionSite = injectionSite(:,:,end:-1:1);
                injectionZ=115-injectionZ;
                
                con=reshape(injectionImg/injectionVolume,size(ANOGD,1)*size(ANOGD,2)*size(ANOGD,3),1);
                save(sprintf('storage\\id_connectivity\\%d',injCount),'con');

                injectedIDs=[injectedIDs,injCount];
                injectedIndizesToID(size(injectedIndizesToID,1)+1,:)=[index(injectionX,injectionY,injectionZ),injCount];
                disp(sprintf('      InjectionSite size in voxel: %d',sum(injectionSite(:)>=1.0)))
                for i = index(injectionSite>=1.0)'
                    injectedIndizesToID(size(injectedIndizesToID,1)+1,:)=[index(indexX(i),indexY(i),indexZ(i)),injCount];
                end

                injCount=injCount+1;

           
            else
                disp(sprintf('There seems to be an error with %d',id))
            end

        end
    end

    save('storage\\injectedIndizes.mat','injectedIDs','')
    save('storage\\injectedIndizes.mat','injectedIndizesToID','-append')

    disp('Computation complete!')
```
