## Supplementary Data 1 for "BrainTACO: An Explorable Multi-Scale Multi-Modal Brain Transcriptomic And Connectivity Data Resource": get_gene_expression_energy.m

```
% Copyright (C) 2022 VRVis.
% All rights reserved.
% Contact: VRVis Forschungs-GmbH
% 
% Redistribution and use in source and binary forms, with or without
% modification, are permitted provided that the following conditions are met:
% 1. Redistributions of source code must retain the above copyright
%    notice, this list of conditions and the following disclaimer.
% 2. Redistributions in binary form must reproduce the above copyright
%    notice, this list of conditions and the following disclaimer in the
%    documentation and/or other materials provided with the distribution.
% 3. All advertising materials mentioning features or use of this software
%    must display the following acknowledgement:
%    This product includes software developed by the VRVis Forschungs-GmbH.
% 4. Neither the name of the VRVis Forschungs-GmbH nor the
%    names of its contributors may be used to endorse or promote products
%    derived from this software without specific prior written permission.
%
% This function downloads gene expression energy from the Allen Mouse Brain Atlas
% and stores it as amira files

	mkdir('storage')
    mkdir('storage/expressions_energy')
    mkdir('storage/expressions_energy_amira')
	mkdir('storage/expressionxml')

  
    %download Allen Brain Atlas hierarchical ontology file
    urlwrite('http://api.brain-map.org/api/v2/structure_graph_download/1.json','storage/ontology.json')
    
    %download a list of all entrez-ids of all genes available from the
    %Allen brain atlas
    disp('Download all available Entrez-IDs from brain-map.org')
    %urlwrite('http://api.brain-map.org/api/v2/data/query.xml?num_rows=30000&only=genes.entrez_id&criteria=model::Gene,rma::criteria,products[abbreviation$eq%27Mouse%27]','storage/allEntrezInDB.xml');
    root = xmlread('storage/allEntrezInDB.xml');

    elems = root.getElementsByTagName('entrez-id');
    entrezAll = {};

    for i = 0:(elems.getLength-1)
        entrezAll{i+1} =char(elems.item(i).getTextContent);
    end

    %generate an index for the entrez ids, that can be randomized to
    %ensure, that random genes are picked from the database
    entrezIndizes=[];
    for i = 1:length(entrezAll)
        if(length(entrezAll{i})>0)
            entrezIndizes=[entrezIndizes,i];
        end
    end

    disp(sprintf('%d Entrez-IDs downloaded!',length(entrezIndizes)))
    entrezIndizes=entrezIndizes(randperm(length(entrezIndizes))); %randomization of the index
    actEntrezIndizes=0;

    disp('Download 100 micron annotations from brain-map.org')

    %download the annotation of the Allen Mouse Common Coordinate Framework
    %in this case, only because we need the dimension-size of the space
    %urlwrite('http://download.alleninstitute.org/informatics-archive/current-release/mouse_ccf/annotation/ccf_2015/annotation_100.nrrd','storage/annotation_100.nrrd');
    [newANOGD, metaDMASK] = nrrdread('storage/annotation_100.nrrd');
    ANOGD=permute(newANOGD,[2 1 3]);

    disp('Complete!')    
    amountOfFiles=size(entrezIndizes,2)
    for actFile = 1:amountOfFiles

        disp(sprintf('Download gene for file %d',actFile))

        expressionIndex=[]; %indizes of expressionMatrix, which rows already contain downloaded data
        expressionMatrix =(zeros(size(ANOGD,1)/2*size(ANOGD,2)/2*size(ANOGD,3)/2,1));

 

            actEntrezIndizes=actEntrezIndizes+1;
            entrez=entrezAll{entrezIndizes(actEntrezIndizes)};
	
	  if(not(exist(strcat('storage/expressionxml/',entrez,'.xml'), 'file')))
            urlwrite(strcat('http://api.brain-map.org/api/v2/data/query.xml?criteria=model::SectionDataSet,rma::criteria,[failed$eq%27false%27],products[abbreviation$eq%27Mouse%27],genes[entrez_id$eq%27',entrez,'%27]'), strcat('storage/expressionxml/',entrez,'.xml'));
		end
            root = xmlread(strcat('storage/expressionxml/',entrez,'.xml'));

            if str2double(root.item(0).getAttribute('num_rows'))<=0

                disp(sprintf('Could not find %s',entrez))
            else

                ids=[];
                referenceSpace=[];
                try
                    for j=1:2:100
                        selectionDataSet = root.item(0).item(0).getChildNodes.item(j);
                        [ attribute, childnodes ] = goToChildNode( selectionDataSet,'id',0 );
                        id = str2num(attribute);
                        ids=[ids,id];
                        [ attribute, childnodes ] = goToChildNode( selectionDataSet,'reference-space-id',0 );
                        rid = str2num(attribute);
                        referenceSpace=[referenceSpace,rid];
                    end
                catch err
                end

                expressionMatrixCor =sparse(size(ANOGD,1)/2*size(ANOGD,2)/2*size(ANOGD,3)/2,1);
                expressionMatrixSag =sparse(size(ANOGD,1)/2*size(ANOGD,2)/2*size(ANOGD,3)/2,1);
                amountCor=sparse(size(ANOGD,1)/2*size(ANOGD,2)/2*size(ANOGD,3)/2,1);
                amountSag=sparse(size(ANOGD,1)/2*size(ANOGD,2)/2*size(ANOGD,3)/2,1);
                
                disp(sprintf('  Get data for %s (%d)',entrez,size(expressionIndex,2)+1))
                for act=1:size(ids,2)
                    id=ids(act);
                    rid=referenceSpace(act);

                    try


                        if(not(exist(sprintf('storage/expressions_energy/%d/energy.raw',id), 'file')))

                            disp(sprintf('      Download %d/%d images for gene from brain-map.org',act,size(ids,2)))
                            urlwrite(sprintf('http://api.brain-map.org/grid_data/download/%d?include=energy',id), 'storage/temp.zip');
                            unzip('storage/temp.zip',sprintf('storage/expressions_energy/%d',id));
                        else
                            disp(sprintf('      Load %d/%d images for gene from local',act,size(ids,2)))
                        end

                        geneGridSize = [67 41 58];
                        fid = fopen(sprintf('storage/expressions_energy/%d/energy.raw',id), 'r', 'l'  );
                        expression = fread( fid, prod(geneGridSize), 'float' );
                        fclose(fid);
                        expression = reshape( expression, geneGridSize );

                        expression=expression(1:(end-1),1:(end-1),1:(end-1));

                        %mirror
                        expression_mir=expression(1:end,1:end,(end):-1:1);
                        expression(1:end,1:end,(floor(size(expression,3)/2)+1):end)=expression_mir(1:end,1:end,(floor(size(expression_mir,3)/2)+1):end);


                        exp = reshape(expression,size(ANOGD,1)/2*size(ANOGD,2)/2*size(ANOGD,3)/2,1);


                        expBiggerZero=exp;
                        expBiggerZero(expBiggerZero<0)=0;


                        if(rid==9)
                            expressionMatrixCor=(expressionMatrixCor+expBiggerZero);
                            amountCor=amountCor+(exp>=0);
                        end
                        if(rid==10)
                            expressionMatrixSag=(expressionMatrixSag+expBiggerZero);
                            amountSag=amountSag+(exp>=0);
                        end

                    catch err
                        disp(sprintf('            Could not open %s: %s',id,getReport(err)))
                    end
                end

                %this is the average computation if multiplie sagital/coronal
                %volumes are available. It excludes "no data" voxels
                %before the average is computed, the gene expression for
                %every grid point is standardized with the mean and std
                %computed by estimate_Mean_STD_of_GeneExpression
                if(sum(expressionMatrixCor(:)>0) && sum(expressionMatrixSag(:)>0))
                    expressionMatrixCor=(expressionMatrixCor./amountCor);
                    expressionMatrixSag=(expressionMatrixSag./amountSag);

                    isnanCor=isnan(expressionMatrixCor);
                    isnanSag=isnan(expressionMatrixSag);

                    divideBy=(2-(isnanCor+isnanSag));
                    expressionMatrixCor(isnanCor & not(isnanSag))=0;
                    expressionMatrixSag(isnanSag & not(isnanCor))=0;

                    expressionMatrixCor(isnanCor & (isnanSag))=NaN;
                    expressionMatrixSag(isnanSag & (isnanCor))=NaN;

                    expressionMatrix(:,1)=(expressionMatrixCor+expressionMatrixSag)./divideBy;
                    expressionIndex= [expressionIndex,size(expressionIndex,2)+1];
                else if(sum(expressionMatrixCor(:)>0) && sum(expressionMatrixSag(:)==0))
                        expressionMatrixCor=(expressionMatrixCor./amountCor);
                        expressionMatrix(:,1)=expressionMatrixCor;
                        expressionIndex= [expressionIndex,size(expressionIndex,2)+1];
                    else  if(sum(expressionMatrixCor(:)==0) && sum(expressionMatrixSag(:)>0))

                            expressionMatrixSag=(expressionMatrixSag./amountSag);
                            expressionMatrix(:,1)=expressionMatrixSag;
                            expressionIndex= [expressionIndex,size(expressionIndex,2)+1];
                        end
                    end
                end


            end
            
    
               
        
             
            
           expressionMatrix=log2(expressionMatrix);
            expressionMatrix(expressionMatrix<0)=0;
            expressionMatrix(isnan(expressionMatrix))=0;
 disp(sprintf('            Maxvalue: %f',max(expressionMatrix)))
            
        
			expressionMatrix=expressionMatrix/6;
			expressionMatrix(expressionMatrix>1)=1;
			fid=fopen(sprintf('storage/expressions_energy_amira/%s.am',entrez),'w');
           
			zippedData = dzip(uint8((expressionMatrix)*255));
            fprintf(fid,sprintf('# AmiraMesh 3D BINARY-LITTLE-ENDIAN 2.0\n# CreationDate: Thu Dec 29 10:01:49 2011\ndefine Lattice %d %d %d\nParameters {\nContent "%dx%dx%d byte, uniform coordinates"\nBoundingBox 0.000000 %d 0.000000 %d 0.000000 %d,\nCoordType "uniform"\n}\nLattice { byte Data } @1(HxZip,%d)\n# Data section follows\n@1\n',size(ANOGD,1)/2,size(ANOGD,2)/2,size(ANOGD,3)/2,size(ANOGD,1)/2,size(ANOGD,2)/2,size(ANOGD,3)/2,size(ANOGD,1),size(ANOGD,2),size(ANOGD,3),length(zippedData)));

            fwrite(fid,zippedData,'uint8');
	
           fclose(fid);
      

        disp('Computation complete!')
    end
	
	
```
