## Supplementary Data 1 for "BrainTACO: An Explorable Multi-Scale Multi-Modal Brain Transcriptomic And Connectivity Data Resource": GeneExpressionTestdata.R

```
## ---- dataset
#### In these files, you need to load a data_matrix (in this case a count matrix), as well as meta data samples

#when load a data matrix, this must be already preprocessed and normalized (e.g. via seurat)
data_matrix<-rbind(
  c(0.0,0.0,0.5,0.5,3.5,3.5,0.0,1.5,6.0,1.3),
	c(0.0,0.0,0.0,0.0,0.0,0.0,0.0,0.0,0.0,0.0),
	c(0.0,0.0,0.3,0.6,6.5,1.5,0.5,1.4,0.0,1.2),
	c(0.0,0.0,0.0001,0.0,1.0,3.2,0.0,0.0,0.0,0)
)
rownames(data_matrix)<-c("Prkcd","Snap25","Th","Adat1")
colnames(data_matrix)<-paste0("TestSample",1:10)

                                  
data_matrix<-log2(((exp(data_matrix)-1)/10000)*1000000+1) #from log1p(data_matrix * 10000) to log2(data_matrix * 1000000 + 1)

#here we have several custom brain regions that we map to brain regions in the mouse
#a testregion 1 --> Hypothalamus
#testregion AAAAA (subregion of BBBBB) --> Cerebral cortex
#testregion BBBBB" --> Cerebrum
#a testregion 2 --> Thalamus
#a testregion 3 (subregion of testregion 2) --> Thalamus, sensory-motor cortex related

#the overlap detection in the main code will change "Cerebrum" to "Cerebral nuclei", which is a sibling of Cerebral Cortex, because Cerebral Cortex is a child of Cerebrum
#The overlap detection in the main code will change "Thalamus" to "Thalamus, polymodal association cortex related", which is a sibling of "Thalamus, sensory-motor cortex related", becasue "Thalamus, sensory-motor cortex related" is a child of "Thalamus"


meta_data_samples<-data.frame(sampleID=colnames(data_matrix),
                              brainRegionStudied=c("a testregion 1","a testregion 1","a testregion 1","a testregion 1","testregion AAAAA (subregion of BBBBB)","testregion BBBBB","a testregion 2","a testregion 2","a testregion 2","a testregion 3 (subregion of testregion 2)"),
                              brainRegionStudiedMapped=c("Hypothalamus","Hypothalamus","Hypothalamus","Hypothalamus","Cerebral cortex","Cerebrum","Thalamus","Thalamus","Thalamus, sensory-motor cortex related
","Thalamus, sensory-motor cortex related"),
                              sampleName=colnames(data_matrix),
                              sex=c("female","female","female","female","female","female","male","male","male","male"),
                              ageCategory=rep("N/A",ncol(data_matrix)),
                              strain=rep("N/A",ncol(data_matrix)),
                              phenotype=rep("N/A",ncol(data_matrix)),
                              genotype=rep("N/A",ncol(data_matrix)),
                              celltype_annotation=c("TestCelltypeAnnotation","TestCelltypeAnnotation1","TestCelltypeAnnotation3","TestCelltypeAnnotation3","TestCelltypeAnnotation3","TestCelltypeAnnotation4","TestCelltypeAnnotation1","TestCelltypeAnnotation1","TestCelltypeAnnotation1","TestCelltypeAnnotation2"),
                              extended_annotation=rep("N/A",ncol(data_matrix)),
                              celltype=c("TestCelltype1","TestCelltype1","TestCelltype3","TestCelltype4","TestCelltype5","TestCelltype6","TestCelltype2","TestCelltype2","TestCelltype2","TestCelltype2"),
                              stringsAsFactors=FALSE)

head(meta_data_samples)

datasetJson<-list()
datasetJson[["name"]]<-"GeneExpressionTestdata"
datasetJson[["description"]]<-"GeneExpressionTestdata"
datasetJson[["doi"]]<-""
datasetJson[["licence"]]<-""
datasetJson[["method"]]<-"scRNA-Seq"
datasetJson[["species"]]<-"Mus musculus" #either Homo sapiens or Mus musculus
datasetJson[["valueTransformation"]]<-"log2+1"
datasetJson[["valueUnit"]]<-"CPM (Counts per Millions)"
```
