## Supplementary Data 1 for "BrainTACO: An Explorable Multi-Scale Multi-Modal Brain Transcriptomic And Connectivity Data Resource": mapping_datasets.nb.html

R Notebook for mapping gene expression datasset to reference spaces


Code 

- Show All Code
- Hide All Code
- Download Rmd

### R Notebook for mapping gene expression datasset to reference spaces


```
datasets<-c(
  "GeneExpressionTestdata" #1
)

#This is only used if one calls convert_datasets.R via Rscript to filter datasets (e.g. for batch processing)
datasetArgumentFilter<-as.numeric(commandArgs(trailingOnly=TRUE))
if(length(datasetArgumentFilter)>0){
  datasets<-datasets[datasetArgumentFilter]
}

computeSeurat<-FALSE #set to trueif you want to compute a tsne plot and also store results as seurat object

datasetToRun<-datasets[1] #only replace the index number if to run this. Don't replace "datasetToRun<-datasets", since it is needed to generate the R script in chunk "Purl"

print(paste0("Run dataset:  ",datasetToRun))
```


```
[1] "Run dataset:  GeneExpressionTestdata"
```


```
projectDir<paste0(dirname(rstudioapi::getSourceEditorContext()$path),"/")
```


```
[1] TRUE
```


```
outputDir<-paste0(dirname(rstudioapi::getSourceEditorContext()$path),"/")
workingDir<-paste0(dirname(rstudioapi::getSourceEditorContext()$path),"/")
```


```
setwd(workingDir)

if(sum(!(c("Matrix","openxlsx","R.matlab","rjson","hash","knitr") %in% installed.packages()[,"Package"]))>0){
  install.packages("Matrix", repos = "http://cran.at.r-project.org/")
  install.packages("openxlsx", repos = "http://cran.at.r-project.org/")
  install.packages("R.matlab", repos = "http://cran.at.r-project.org/")
  install.packages("rjson", repos = "http://cran.at.r-project.org/")
  install.packages("hash", repos = "http://cran.at.r-project.org/")
  install.packages("knitr", repos = "http://cran.at.r-project.org/")
}

library(Matrix)
library(openxlsx) #Needed to read xlsm files
library(R.matlab) #Needed to read atlas files
library(rjson) #Needed to read the AHBA ontology
library(hash)
library(knitr)
library(Seurat)
```


```
#creates a script for each dataset to run

if(file.exists(paste0("convert_datasets.Rmd"))){
  unlink("convert_datasets.R")
  purl("convert_datasets.Rmd", output = paste0("convert_datasets.R"),documentation = 2)
  
  #add for loop
  linesOfR = readLines("convert_datasets.R")
  for(lineIndex in 1:length(linesOfR)){
     if(startsWith(linesOfR[lineIndex], "datasetToRun<-datasets")){
        linesOfR[lineIndex]<-"for(datasetToRun in datasets){"
    }
  }
  linesOfR<-c(linesOfR,"}") 
  writeLines(linesOfR, "convert_datasets.R")
}
```


```
unlink(paste0(outputDir,datasetToRun,".log"))
sink(paste0(outputDir,datasetToRun,".log"), append=FALSE, split=TRUE) 
print(paste0("Initialized logging for dataset:  ",datasetToRun))
```


```
[1] "Initialized logging for dataset:  GeneExpressionTestdata"
```


```
atlasRegionsMouseFile<-paste0(workingDir,"//","storage/atlasRegions.mat")
atlasRegionsHumanFile<-paste0(workingDir,"//","storage/atlasRegionsHumanAHRA.mat")

ontologyMouseFile<-paste0(workingDir,"//","storage/ontology.json")
ontologyHumanFile<-paste0(workingDir,"//","storage/ontologyHumanAHRA.json")
```


```
#checks ensemble id from gene symbol for by ensemble.org via bioMart (is only used by dataset specific sub scripts) for mouse
getIDFromSymbolMouse <- function(symbol_Vector){
    library(biomaRt)
    out <- tryCatch(
            {
              gene_name_ensemblid_1<-getBM(
                attributes = c("ensembl_gene_id","external_gene_name"),
                filters = "external_gene_name",
                values = symbol_Vector,
                mart = useMart("ensembl", dataset = "mmusculus_gene_ensembl",host="uswest.ensembl.org"),
              )
              gene_name_ensemblid_2<-getBM(
                attributes = c("ensembl_gene_id","uniprot_gn_symbol"), 
                filters = "uniprot_gn_symbol", 
                values = symbol_Vector, 
                mart = useMart("ensembl", dataset = "mmusculus_gene_ensembl",host="uswest.ensembl.org"),
              )
              
              gene_name_ensemblid_3<-getBM(
                attributes = c("ensembl_gene_id","mgi_symbol"), 
                filters = "mgi_symbol", 
                values = symbol_Vector, 
                mart = useMart("ensembl", dataset = "mmusculus_gene_ensembl",host="uswest.ensembl.org"),
              )
              
               gene_name_ensemblid_4<-getBM(
                attributes = c("ensembl_gene_id","external_synonym"), 
                filters = "external_synonym", 
                values = symbol_Vector, 
                mart = useMart("ensembl", dataset = "mmusculus_gene_ensembl",host="uswest.ensembl.org"),
              )
               
                genes_in_db <- read.csv2(paste0(workingDir,"//","storage/genes.csv"),header=TRUE,stringsAsFactors = FALSE)
                genes_in_db <- genes_in_db[genes_in_db$SPECIES=="Mus musculus",]
            
                gene_name_ensemblid_5 <- cbind(rep("",length(symbol_Vector)),rep("",length(symbol_Vector)))
              
                for(actGeneRow in 1:length(symbol_Vector)){
                  indexFound<-c()
                 
                  if(sum(genes_in_db$SYMBOL==symbol_Vector[actGeneRow])==0){ #r replaces duplicated rownames in rownames with "." -> if that happend in the preprocessing of the data, we can account for this by removing the . and everything after that
                    indexFound<-which(genes_in_db$SYMBOL==strsplit(symbol_Vector[actGeneRow], "\\.")[[1]][1])
                  }else{
                    indexFound<-which(genes_in_db$SYMBOL==symbol_Vector[actGeneRow])
                  }
                  
                  indexFound<-indexFound[!is.na(indexFound)]
                  
                  if(length(indexFound)>0){
                    gene_name_ensemblid_5[actGeneRow,]<-c(genes_in_db$ENSEMBL[indexFound[1]],symbol_Vector[actGeneRow])
                  }
                }   
                gene_name_ensemblid_5<-gene_name_ensemblid_5[gene_name_ensemblid_5[,1]!="" & !is.na(gene_name_ensemblid_5[,1]),]   
             
              gene_name_ensemblid<-base::rbind(as.matrix(gene_name_ensemblid_1),as.matrix(gene_name_ensemblid_2),as.matrix(gene_name_ensemblid_3),as.matrix(gene_name_ensemblid_4),as.matrix(gene_name_ensemblid_5))
              gene_name_ensemblid<-gene_name_ensemblid[!duplicated(gene_name_ensemblid[,2]),]
              
                 
              library("org.Mm.eg.db")
              gene_name_entrezids<-mapIds(org.Mm.eg.db, keys = symbol_Vector, keytype = "SYMBOL", column=c("ENTREZID"))
              gene_name_entrezids<-gene_name_entrezids[!is.na(gene_name_entrezids)]
              gene_name_entrezids<-cbind(gene_name_entrezids,names(gene_name_entrezids))
              
              if(dim(gene_name_entrezids)[1]>dim(gene_name_ensemblid)[1]){
                return(gene_name_entrezids)
              }else{
                 return(gene_name_ensemblid)
              }
              
              return(gene_name_ensemblid)
            },
            error=function(cond) {
                print(paste("Could not reach biomRt server, no matching of symbols to ensemblid possible! Code will work without (matching will performed offline), but offline matching might lead to worse results."))
               
                return(cbind(symbol_Vector,symbol_Vector))
            }
      )    
      return(out)
}

#checks ensemble id from gene symbol for by ensemble.org via bioMart (is only used by dataset specific sub scripts) for human
getIDFromSymbolHuman <- function(symbol_Vector){
  library(biomaRt)
    out <- tryCatch(
            {
              gene_name_ensemblid_1<-getBM(
                attributes = c("ensembl_gene_id","external_gene_name"),
                filters = "external_gene_name",
                values = symbol_Vector,
                mart = useMart("ensembl", dataset = "hsapiens_gene_ensembl",host="uswest.ensembl.org"),
              )
              gene_name_ensemblid_2<-getBM(
                attributes = c("ensembl_gene_id","uniprot_gn_symbol"), 
                filters = "uniprot_gn_symbol", 
                values = symbol_Vector, 
                mart = useMart("ensembl", dataset = "hsapiens_gene_ensembl",host="uswest.ensembl.org"),
              )
              
              gene_name_ensemblid_3<-getBM(
                attributes = c("ensembl_gene_id","hgnc_symbol"), 
                filters = "hgnc_symbol", 
                values = symbol_Vector, 
                mart = useMart("ensembl", dataset = "hsapiens_gene_ensembl",host="uswest.ensembl.org"),
              )
              
              gene_name_ensemblid_4<-getBM(
                attributes = c("ensembl_gene_id","external_synonym"), 
                filters = "external_synonym", 
                values = symbol_Vector, 
                mart = useMart("ensembl", dataset = "hsapiens_gene_ensembl",host="uswest.ensembl.org"),
              )
              
                genes_in_db <- read.csv2(paste0(workingDir,"//","storage/genes.csv"),header=TRUE,stringsAsFactors = FALSE)
                genes_in_db <- genes_in_db[genes_in_db$SPECIES=="Homo sapiens",]
            
                gene_name_ensemblid_5 <- cbind(rep("",length(symbol_Vector)),rep("",length(symbol_Vector)))
              
                for(actGeneRow in 1:length(symbol_Vector)){
                  indexFound<-c()
                 
                  if(sum(genes_in_db$SYMBOL==symbol_Vector[actGeneRow])==0){ #r replaces duplicated rownames in rownames with "." -> if that happend in the preprocessing of the data, we can account for this by removing the . and everything after that
                    indexFound<-which(genes_in_db$SYMBOL==strsplit(symbol_Vector[actGeneRow], "\\.")[[1]][1])
                  }else{
                    indexFound<-which(genes_in_db$SYMBOL==symbol_Vector[actGeneRow])
                  }
                  
                  indexFound<-indexFound[!is.na(indexFound)]
                  
                  if(length(indexFound)>0){
                    gene_name_ensemblid_5[actGeneRow,]<-c(genes_in_db$ENSEMBL[indexFound[1]],symbol_Vector[actGeneRow])
                  }
                }   
                gene_name_ensemblid_5<-gene_name_ensemblid_5[gene_name_ensemblid_5[,1]!="" & !is.na(gene_name_ensemblid_5[,1]),]   
             
              
              gene_name_ensemblid<-base::rbind(as.matrix(gene_name_ensemblid_1),as.matrix(gene_name_ensemblid_2),as.matrix(gene_name_ensemblid_3),as.matrix(gene_name_ensemblid_4),as.matrix(gene_name_ensemblid_5))
              gene_name_ensemblid<-gene_name_ensemblid[!duplicated(gene_name_ensemblid[,2]),]
              
              library("org.Hs.eg.db")
              gene_name_entrezids<-mapIds(org.Hs.eg.db, keys = symbol_Vector, keytype = "SYMBOL", column=c("ENTREZID"))
              gene_name_entrezids<-gene_name_entrezids[!is.na(gene_name_entrezids)]
              gene_name_entrezids<-cbind(gene_name_entrezids,names(gene_name_entrezids))
              
              if(dim(gene_name_entrezids)[1]>dim(gene_name_ensemblid)[1]){
                return(gene_name_entrezids)
              }else{
                 return(gene_name_ensemblid)
              }
          },
            error=function(cond) {
                print(paste("Could not reach biomRt server, no matching of symbols to ensemblid possible! Code will work without (matching will performed offline), but offline matching might lead to worse results."))
               
                return(cbind(symbol_Vector,symbol_Vector))
            }
      )    
      return(out)
  
}
```


```
  data_matrix<-NULL
```


```
source(paste0('dataset_chunks/',datasetToRun,".R"))
```


```
print(paste0("Data matrix has ",nrow(data_matrix)," genes matched to gene database and ",ncol(data_matrix)," samples"))
```


```
[1] "Data matrix has 4 genes matched to gene database and 10 samples"
```


```
if(is.null(colnames(data_matrix))){
  error("ERROR, data_matrix does not have colnames, should be sample/measurement ids!")
}
if(is.null(rownames(data_matrix))){
  error("ERROR, data_matrix does not have rownames, should be genes ids!")
}
if(!is.numeric(data_matrix[1,1])){
  error("ERROR, data_matrix does not have numeric elements!")
}
```


```
  datasetJson[["name"]]<-gsub(" ","_",datasetJson[["name"]])
  datasetJson[["maximumValue"]]<-"0"

  if(datasetJson[["species"]]=="Homo sapiens" || datasetJson[["species"]]=="Macaca mulatta"){
    datasetJson[["brainRegionParcellation"]]<-"Human AHRA Hierarchy"
  }
  if(datasetJson[["species"]]=="Mus musculus" || datasetJson[["species"]]=="Rattus norvegicus"){
    datasetJson[["brainRegionParcellation"]]<-"AMBA 100 micron hierarchy"
  }
  
  replaceSamplesWithMeasurements<-data.frame() #in case of brain activity data
  
  #if idType is not set, we assume it is gene expression, so we determine it based on the data_matrix row names
  #NOTE: DON'T USE SYMBOL, BRAINTRAWLER IS BAD A MATCHING THOSE
  if(is.null(datasetJson[["idType"]])){
    if(suppressWarnings(is.na(as.numeric(rownames(data_matrix)[1])))){  #either entrezid or ensemblid
      if(grepl("EN",rownames(data_matrix)[1])){
        datasetJson[["idType"]]<-"ensemblid"
        print("Genes are identfied by ensemblid")
      }else{
        datasetJson[["idType"]]<-"symbol"
        print("Genes are identfied by symbol")
      }
    }else{
      datasetJson[["idType"]]<-"entrezid"
      print("Genes are identfied by entrezid")
    }
  }
```


```
[1] "Genes are identfied by symbol"
```


```
if(is.null(data_matrix)){
  break
}
```


```
  #this checks if rna rows match with meta data file
  if( sum(is.element(colnames(data_matrix),meta_data_samples$sampleID))<
      sum(is.element(colnames(data_matrix),gsub("-",".",meta_data_samples$sampleID)))
  ){
    meta_data_samples$sampleID<-gsub("-",".",meta_data_samples$sampleID)
  } 
  
  if(length(intersect(colnames(data_matrix),meta_data_samples$sampleID))!=length(colnames(data_matrix))){
    print(paste0("Warning: could only match ",length(intersect(colnames(data_matrix),meta_data_samples$sampleID))/length(colnames(data_matrix))*100,"% data_matrix columns with sampleIDs!"))
    #setdiff(colnames(data_matrix),meta_data_samples$sampleID)[1]
  }
  
  #this checks if rna rows match with meta data file
  if(sum(colnames(data_matrix)==meta_data_samples$sampleID)!=length(colnames(data_matrix))){
    print(paste0("Warning: only ",sum(colnames(data_matrix)==meta_data_samples$sampleID)/length(colnames(data_matrix))*100,"% data_matrix columns allign with sampleIDs! Rearrange data_matrix"))
    
    print(paste0("RNA data dimension before sample alignment: ",dim(data_matrix)[1],"x",dim(data_matrix)[2]))
      print(paste0("Amount of samples in meta data: ",length(meta_data_samples$sampleID)))
    
    # if(length(meta_data_samples$sampleID)>dim(data_matrix)[2]){
    #   print("Warning, more sample information than in RNA data, fixing length...")
        meta_data_samples<-meta_data_samples[is.element(meta_data_samples$sampleID,colnames(data_matrix)),]
        print(paste0("New amount of samples in meta data: ",length(meta_data_samples$sampleID)))
    # }
    
    print(paste0("Example for RNA data sample: ",colnames(data_matrix)[1]))
      print(paste0("Example for meta data sample: ",meta_data_samples$sampleID[1]))
    
    data_matrix<-data_matrix[,meta_data_samples$sampleID]
    print(paste0("... and after sample alignment: ",dim(data_matrix)[1],"x",dim(data_matrix)[2]))
    
    print(paste0("Now ",sum(colnames(data_matrix)==meta_data_samples$sampleID)/length(colnames(data_matrix))*100,"% data_matrix columns allign with sampleIDs!"))
    
  }
```


```
ontology<-c() #hierarchical ontology
atlasRegions<-c() #the reference space (3D array, with region ids from ontology)

if(datasetJson[["species"]]=="Homo sapiens" || datasetJson[["species"]]=="Macaca mulatta"){
  atlasRegions<-readMat(atlasRegionsHumanFile)$atlasRegions #Atlas IDs of voxels
  ontology <- fromJSON(file=ontologyHumanFile, method='C')   #get ontology
}

if(datasetJson[["species"]]=="Mus musculus" || datasetJson[["species"]]=="Rattus norvegicus"){
  atlasRegions<-readMat(atlasRegionsMouseFile)$atlasRegions #Atlas IDs of voxels
  ontology <- fromJSON(file=ontologyMouseFile, method='C')   #get ontology
}
```


```
print("Generate mapping from brain regions to voxels")
```


```
[1] "Generate mapping from brain regions to voxels"
```


```
#remove unintentional "\n"
meta_data_samples$brainRegionStudiedMapped<-sapply(meta_data_samples$brainRegionStudiedMapped,function(x){
  gsub("\\n", "", x)
})
#remove unintentional spaces before name"
meta_data_samples$brainRegionStudiedMapped<-sapply(meta_data_samples$brainRegionStudiedMapped,function(x){
  gsub("; ", ";", x)
}) 

regions_to_atlasIDs<-data.frame(brainRegions=unique(meta_data_samples$brainRegionStudiedMapped),
                          atlasID=rep("",length(unique(meta_data_samples$brainRegionStudiedMapped))),
                          stringsAsFactors=FALSE)

#uses the mapping of brainRegionStudiedMapped to map the regions to the atlasIDs
#It will automatically add L_ and R_ split mapping if not indicated in the mapping (both hemispheres)
for(actRow in 1:nrow(regions_to_atlasIDs)){
  newString<-""
  prefix<-""
  for(splitBrainRegion in strsplit(regions_to_atlasIDs$brainRegions[actRow],";")[[1]]){
    ontologyInformationOfRegion<-getIDByName(ontology$msg[[1]],splitBrainRegion)
    if(is.null(ontologyInformationOfRegion)){
      searchstring<-strsplit(splitBrainRegion,'\\ \\(')[[1]][1]
    }else{
      searchstring<-splitBrainRegion
    }
    
    if(!is.na(searchstring)){
      ontologyInformationOfRegion<-getIDByName(ontology$msg[[1]],searchstring)
      if(!is.null(ontologyInformationOfRegion)){
        if(regions_to_atlasIDs$atlasID[actRow]!=""){
          regions_to_atlasIDs$atlasID[actRow]<-paste0(regions_to_atlasIDs$atlasID[actRow],";",ontologyInformationOfRegion$id)
        }else{
          regions_to_atlasIDs$atlasID[actRow]<-paste(ontologyInformationOfRegion$id)
        }
        newString<-paste0(newString,prefix,paste0("L_",ontologyInformationOfRegion$name,";R_",ontologyInformationOfRegion$name))
        prefix<-";"
        print(paste0("MAPPED: ",searchstring," TO ",ontologyInformationOfRegion$name, " ID: ",ontologyInformationOfRegion$id))
      }else{
        splitLeftRight<-strsplit(splitBrainRegion,'L_')[[1]]
        if(length(splitLeftRight)==2){
          prefixLeftRight<-"L_"
        }else{
          splitLeftRight<-strsplit(splitBrainRegion,'R_')[[1]]
          if(length(splitLeftRight)==2){
            prefixLeftRight<-"R_"
          }
        }
        
        if(length(splitLeftRight)==2){
          ontologyInformationOfRegion<-getIDByName(ontology$msg[[1]],splitLeftRight[2])
          if(!is.null(ontologyInformationOfRegion)){
            if(regions_to_atlasIDs$atlasID[actRow]!=""){
              regions_to_atlasIDs$atlasID[actRow]<-paste0(regions_to_atlasIDs$atlasID[actRow],";",ontologyInformationOfRegion$id)
            }else{
              regions_to_atlasIDs$atlasID[actRow]<-paste(ontologyInformationOfRegion$id)
            }
            newString<-paste0(newString,prefix,paste0(prefixLeftRight,ontologyInformationOfRegion$name))
            prefix<-";"
            print(paste0("MAPPED: ",splitBrainRegion," TO ",prefixLeftRight,ontologyInformationOfRegion$name, " ID: ",ontologyInformationOfRegion$id))
          }else{
            print(paste0("NO_MAPPING: ",splitBrainRegion," TO ",ontologyInformationOfRegion$name, " ID: NULL"))
          }
          
        }else{
          print(paste0("NO_MAPPING: ",searchstring," TO ",ontologyInformationOfRegion$name, " ID: NULL"))
        }
      }
    }else{
      print(paste0("NO_MAPPING: ",searchstring," TO NULL ID: NULL"))
    }
  }
  if(nchar(newString)>0){
    meta_data_samples$brainRegionStudiedMapped[meta_data_samples$brainRegionStudiedMapped==regions_to_atlasIDs$brainRegions[actRow]]<-newString
    datasetJson[["samples"]]$brainRegionStudiedMapped[datasetJson[["samples"]]$brainRegionStudiedMapped==regions_to_atlasIDs$brainRegions[actRow]]<-newString
    regions_to_atlasIDs$brainRegions[actRow]<-newString
  }
  
}
```


```
[1] "MAPPED: Hypothalamus TO Hypothalamus ID: 1097"
[1] "MAPPED: Cerebral cortex TO Cerebral cortex ID: 688"
[1] "MAPPED: Cerebrum TO Cerebrum ID: 567"
[1] "MAPPED: Thalamus TO Thalamus ID: 549"
[1] "MAPPED: Thalamus, sensory-motor cortex related TO Thalamus, sensory-motor cortex related ID: 864"
```


```
#Remove regions without mapping
regions_to_atlasIDs<-regions_to_atlasIDs[regions_to_atlasIDs$atlasID!="",]

brainRegionSave<-hash()
#add the voxel level representation of brain regions to a hashmap, so we don't have to generate them every time they are needed
for(brainRegionRow in 1:nrow(regions_to_atlasIDs)){
  isFirst<-TRUE
  for(splitRegionID in strsplit(regions_to_atlasIDs$atlasID[brainRegionRow],";")[[1]]){
    if(isFirst){
      brainRegionSave[[paste(regions_to_atlasIDs$atlasID[brainRegionRow])]]<-getAtlasRegionsOfID(atlasRegions,as.numeric(splitRegionID))>0 
      isFirst<-FALSE
    }else{
      brainRegionSave[[paste(regions_to_atlasIDs$atlasID[brainRegionRow])]]<- brainRegionSave[[paste(regions_to_atlasIDs$atlasID[brainRegionRow])]] | getAtlasRegionsOfID(atlasRegions,as.numeric(splitRegionID))>0 
    }
  }
}

print("")
```


```
[1] ""
```


```
print("Check for overlapping regions:")
```


```
[1] "Check for overlapping regions:"
```


```
#Check for overlapping regions.
#It will automatically remove overlaps in this way:
#If RegionA is the parent of RegionA_A, RegionA_B and RegionA_C, and there are is a sample with RegionA and one with RegionA_A. 
#In this case, it will always thake the most detailed mapping (down the hierarchy, i.e. leaves would be the most accurate ones)
#Since RegionA_C is already associated to a sample, RegionA can not cover it. Hence, RegionA will be cahnged to "RegionA_A;RegionA_B"
for(brainRegionRow in 1:nrow(regions_to_atlasIDs)){
  hasSubregions<-FALSE
  mergedText<-""
  prefix<-""
  for(otherRegionRow in 1:nrow(regions_to_atlasIDs)){
    otherRegionIsSubregion<-FALSE
   
    for(splitRegionIDRow in strsplit(regions_to_atlasIDs$atlasID[brainRegionRow],";")[[1]]){
      for(splitRegionIDOtherRow in strsplit(regions_to_atlasIDs$atlasID[otherRegionRow],";")[[1]]){
        #it is a subregion if it is a direct subregion
        otherRegionIsSubregion <- otherRegionIsSubregion | isSubregion(ontology$msg[[1]],as.numeric(splitRegionIDRow),as.numeric(splitRegionIDOtherRow))
        #it is also a subregion brainRegionRow contains of more regions than otherRegionRow and the same region
         otherRegionIsSubregion <- otherRegionIsSubregion | (length(strsplit(regions_to_atlasIDs$atlasID[brainRegionRow],";")[[1]])>length(strsplit(regions_to_atlasIDs$atlasID[otherRegionRow],";")[[1]]) & as.numeric(splitRegionIDRow)==as.numeric(splitRegionIDOtherRow))
      }
    }
    
    if(otherRegionIsSubregion){ 
      brainRegionSave[[paste(regions_to_atlasIDs$atlasID[brainRegionRow])]]<-(brainRegionSave[[paste(regions_to_atlasIDs$atlasID[brainRegionRow])]]-brainRegionSave[[paste(regions_to_atlasIDs$atlasID[otherRegionRow])]])>0
      mergedText<-paste0(mergedText,prefix,paste0(regions_to_atlasIDs$brainRegions[otherRegionRow]))
      prefix<-","
      hasSubregions<-TRUE         
      
    }
  }
  
  #change mapping if there are subregions. e.g. If you have a region mapped to Striatum, and one to Lateral septal complex (which is a subregion of the Striatum), then Striatum will be mapped to all subregions of Striatum excet the Lateral septal complex
  if(hasSubregions){
    if(nchar(mergedText)>40){
        print(paste0("CHANGE MAPPING OF ",regions_to_atlasIDs$brainRegions[brainRegionRow]," BECAUSE OF SUBREGION(s): ",strtrim(mergedText, 40),"..."))
    }else{
      print(paste0("CHANGE MAPPING OF ",regions_to_atlasIDs$brainRegions[brainRegionRow]," BECAUSE OF SUBREGION(s): ",mergedText))
    }
  
    
    restOfBrainRegionIDs<-unique(atlasRegions[brainRegionSave[[paste(regions_to_atlasIDs$atlasID[brainRegionRow])]]]) #this will be iteratively reduced to contain only ids that do not cover otherRegionRow
    idsToCheck<-restOfBrainRegionIDs
    
    while(length(idsToCheck)>0){ #this code will go through the hierarchy to find the least amount of regions discribing "the rest of the brain"
      #print(paste0("idsToCheck length: ",length(idsToCheck)," length of unique restOfBrainRegionIDs: ",length(unique(restOfBrainRegionIDs))))
   
      idToCheck<-idsToCheck[1]

      ontologyInformationOfRegionIDToCheck<-getAtlasRegionByID(ontology$msg[[1]],idToCheck)
      if(!is.null(ontologyInformationOfRegionIDToCheck$parent_structure_id)){
        hasSubregions<-FALSE
        for(otherRegionRow in 1:nrow(regions_to_atlasIDs)){#If the other regions are not a subregion of the parent of the id to check, replace idTo check with parent
          for(splitRegionIDOtherRow in strsplit(regions_to_atlasIDs$atlasID[otherRegionRow],";")[[1]]){
            hasSubregions <- hasSubregions | (isSubregion(ontology$msg[[1]],ontologyInformationOfRegionIDToCheck$parent_structure_id,as.numeric(splitRegionIDOtherRow)) || as.numeric(splitRegionIDOtherRow)==ontologyInformationOfRegionIDToCheck$parent_structure_id)
          }
        }
        if(hasSubregions){
          idsToCheck<-idsToCheck[-1]
        }else{
          idsToCheck[idsToCheck==idToCheck]<-ontologyInformationOfRegionIDToCheck$parent_structure_id
          restOfBrainRegionIDs[restOfBrainRegionIDs==idToCheck]<-ontologyInformationOfRegionIDToCheck$parent_structure_id
        }
      }else{
        #in this case, the id to check must be root
        idsToCheck<-idsToCheck[-1]
      }
    }
    restOfBrainRegionIDs<-restOfBrainRegionIDs[!is.element(restOfBrainRegionIDs,regions_to_atlasIDs$atlasID[brainRegionRow])] #could be that the act brain region (brainRegionRow) has als some undefined parts (voxels without a subregion, i.e. some voxels directly map to voxel) so we remove it since we can't map this to the coordinates to region definition file
    
    prefix<-""
    mergedText<-""
    mergedID<-""
    for(restOfBrainRegionID in unique(restOfBrainRegionIDs)){
      ontologyInformationOfRegionRestOfBrainRegionID<-getAtlasRegionByID(ontology$msg[[1]],restOfBrainRegionID)
      if(grepl("L_", regions_to_atlasIDs$brainRegions[brainRegionRow]) && grepl("R_", regions_to_atlasIDs$brainRegions[brainRegionRow])){
        mergedText<-paste0(mergedText,prefix,paste0("L_",ontologyInformationOfRegionRestOfBrainRegionID$name,";R_",ontologyInformationOfRegionRestOfBrainRegionID$name))
        mergedID<-paste0(mergedID,prefix,ontologyInformationOfRegionRestOfBrainRegionID$id)
      }else{
        if(grepl("L_", regions_to_atlasIDs$brainRegions[brainRegionRow])){
          mergedText<-paste0(mergedText,prefix,paste0("L_",ontologyInformationOfRegionRestOfBrainRegionID$name))
          mergedID<-paste0(mergedID,prefix,ontologyInformationOfRegionRestOfBrainRegionID$id)
        }else{
          if(grepl("R_", regions_to_atlasIDs$brainRegions[brainRegionRow])){
            mergedText<-paste0(mergedText,prefix,paste0("R_",ontologyInformationOfRegionRestOfBrainRegionID$name))
            mergedID<-paste0(mergedID,prefix,ontologyInformationOfRegionRestOfBrainRegionID$id)
          }
        }
      }
      
      prefix<-";"
    }
    
    if(mergedText==regions_to_atlasIDs$brainRegions[brainRegionRow] || mergedText==""){
      print(paste0("-->Region could not be mapped since there are no regions left that are not covered by subregions. Set ",sum(meta_data_samples$brainRegionStudiedMapped==regions_to_atlasIDs$brainRegions[brainRegionRow])," samples to empty region"))
      meta_data_samples$brainRegionStudiedMapped[meta_data_samples$brainRegionStudiedMapped==regions_to_atlasIDs$brainRegions[brainRegionRow]]<-""
    }else{
      print(paste0("TO: ",mergedText))
      meta_data_samples$brainRegionStudiedMapped[meta_data_samples$brainRegionStudiedMapped==regions_to_atlasIDs$brainRegions[brainRegionRow]]<-mergedText
      regions_to_atlasIDs$brainRegions[brainRegionRow]<-mergedText
      regions_to_atlasIDs$atlasID[brainRegionRow]<-mergedID
    }
  }
}
```


```
[1] "CHANGE MAPPING OF L_Cerebrum;R_Cerebrum BECAUSE OF SUBREGION(s): L_Cerebral cortex;R_Cerebral cortex"
[1] "TO: L_Cerebral nuclei;R_Cerebral nuclei"
[1] "CHANGE MAPPING OF L_Thalamus;R_Thalamus BECAUSE OF SUBREGION(s): L_Thalamus, sensory-motor cortex related..."
[1] "TO: L_Thalamus, polymodal association cortex related;R_Thalamus, polymodal association cortex related"
```


```
for(brainRegionRow in 1:nrow(regions_to_atlasIDs)){
  #If new brain region combination have been created, adapt brianRegionSave
  if(is.null(brainRegionSave[[paste(regions_to_atlasIDs$atlasID[brainRegionRow])]])){
    isFirst<-TRUE
    for(splitRegionID in strsplit(regions_to_atlasIDs$atlasID[brainRegionRow],";")[[1]]){
      if(isFirst){
        brainRegionSave[[paste(regions_to_atlasIDs$atlasID[brainRegionRow])]]<-getAtlasRegionsOfID(atlasRegions,as.numeric(splitRegionID))>0 
        isFirst<-FALSE
      }else{
        brainRegionSave[[paste(regions_to_atlasIDs$atlasID[brainRegionRow])]]<- brainRegionSave[[paste(regions_to_atlasIDs$atlasID[brainRegionRow])]] | getAtlasRegionsOfID(atlasRegions,as.numeric(splitRegionID))>0 
      }
    }
  }
  
  #Adapt for hemispheres
  if(regions_to_atlasIDs$brainRegions[[brainRegionRow]]!=""){
      hemisphere<-getHemisphere(regions_to_atlasIDs$brainRegions[[brainRegionRow]])
      if(hemisphere<3){
        if(is.null(brainRegionSave[[paste0(regions_to_atlasIDs$atlasID[brainRegionRow],"_",hemisphere)]]) && sum(brainRegionSave[[paste(regions_to_atlasIDs$atlasID[brainRegionRow])]])>0){
          hemisphereData<-getAtlasRegionsHemisphere(brainRegionSave[[paste(regions_to_atlasIDs$atlasID[brainRegionRow])]],hemisphere)>0
          regions_to_atlasIDs$atlasID[brainRegionRow]<-paste0(regions_to_atlasIDs$atlasID[brainRegionRow],"_",hemisphere)
          brainRegionSave[[paste0(regions_to_atlasIDs$atlasID[brainRegionRow])]]<-hemisphereData
        }
      }
    }
}

meta_data_samples$brainRegionStudiedMapped[!is.element(meta_data_samples$brainRegionStudiedMapped,regions_to_atlasIDs$brainRegions)]<-"" #set mapping to "" if it could not be mapped

print(paste0("",sum(meta_data_samples$brainRegionStudiedMapped!=""),"/",length(meta_data_samples$brainRegionStudiedMapped)," samples could be mapped to the reference space"))
```


```
[1] "10/10 samples could be mapped to the reference space"
```


```
regionsThatHaveNoVoxelLevelRepresentation<-regions_to_atlasIDs$brainRegions[sapply(1:nrow(regions_to_atlasIDs),function(x){
  sum(brainRegionSave[[paste(regions_to_atlasIDs$atlasID[x])]])==0 
})]

if(length(regionsThatHaveNoVoxelLevelRepresentation)>0){
  print("WARNING: The following regions have no voxel level representation in the reference space and will be removed:")
  for(actRegion in regionsThatHaveNoVoxelLevelRepresentation){
    print(paste0(actRegion))
  }
}

#remove regions without mapping
regions_to_atlasIDs<-regions_to_atlasIDs[sapply(1:nrow(regions_to_atlasIDs),function(x){
  sum(brainRegionSave[[paste(regions_to_atlasIDs$atlasID[x])]])>0 && sum(regions_to_atlasIDs$brainRegions[x]==meta_data_samples$brainRegionStudiedMapped)>0
}),]


datasetJson[["samples"]]<-meta_data_samples


if(length(unique(regions_to_atlasIDs$atlasID))!=length(regions_to_atlasIDs$atlasID)){
  print("WARNING, non-unique atlas IDs, every atlas ID should be here only once!!!")
}
```


```
#creating a region to coordinates file which maps brain regions to coordinates in the reference space (needed for spatial indexing)
  print("Define output data and create regions to coordinates file")
```


```
[1] "Define output data and create regions to coordinates file"
```


```
  suppressWarnings(dir.create(paste0(outputDir,"spatialdata")))
  suppressWarnings(dir.create(paste0(outputDir,"spatialdata/datasets")))
  suppressWarnings(dir.create(paste0(outputDir,"input_data")))
  suppressWarnings(dir.create(paste0(outputDir,"input_data/datasets")))

  #output data json and outputData need the same name!!!
  outputJson<-paste0(outputDir,"input_data/datasets/",datasetJson[["name"]],".json")
  outputData<-paste0(outputDir,"spatialdata/datasets/",datasetJson[["name"]])

  
  suppressWarnings(dir.create(outputData))

  coordinatesRegion<-c()
  coordinatesRegion<-matrix(0,nrow=length(unlist(atlasRegions)),ncol=4)
  amountOfVoxels<-0
  for(actRegionRow in 1:nrow(regions_to_atlasIDs)){
    brainRegion<-brainRegionSave[[paste(regions_to_atlasIDs$atlasID[actRegionRow])]]
    for(x in 1:dim(brainRegion)[1]){
      for(y in 1:dim(brainRegion)[2]){
        for(z in 1:dim(brainRegion)[3]){
          if(brainRegion[x,y,z]>0){
            coordinatesRegion[amountOfVoxels+1,1]<-x
            coordinatesRegion[amountOfVoxels+1,2]<-y
            coordinatesRegion[amountOfVoxels+1,3]<-z
            coordinatesRegion[amountOfVoxels+1,4]<-actRegionRow-1
            amountOfVoxels<-amountOfVoxels+1
          }
        }
      }  
    }
  }
  coordinatesRegion<-coordinatesRegion[1:amountOfVoxels,]
  write.table(coordinatesRegion,paste0(outputData,"/coordinates_to_region_index.csv"),col.names=FALSE,row.names=FALSE,dec=".",sep=",")
```


```
  print("Create count matrix and import data")
```


```
[1] "Create count matrix and import data"
```


```
sample_information <- meta_data_samples[,colnames(meta_data_samples)!="sampleName"]
sample_information[is.na(sample_information)] <- "N/A"

  sample_information$region_id <- unlist(sapply(meta_data_samples$brainRegionStudiedMapped,function(x){
    if(length(regions_to_atlasIDs$brainRegions)>1){
      retVal <- ((1:length(regions_to_atlasIDs$brainRegions))-1)[regions_to_atlasIDs$brainRegions==x]
      if(length(retVal)==1){
        return(retVal)
      }else{
        return(NA)
      }
    }else{
      return(0)
    }
  }),use.names = FALSE)
  
  if(sum(is.na(sample_information$region_id))>0){
    print(paste0("Warning: ",sum(is.na(sample_information$region_id))," samples of ",length(sample_information$region_id)," had no region annotation and will be removed!"))
  }

  sample_information_filtered<-sample_information[!is.na(sample_information$region_id),]
  colnames(sample_information_filtered)[1]<-"sampleID"
  write.csv2(sample_information_filtered,paste0(outputData,"/sample_information.csv"),row.names = FALSE)
  
  
  foundGeneInDatabase <- rep(0,nrow(data_matrix))
  pb <- txtProgressBar(min=0, max=nrow(data_matrix), initial=0,style=3)
```


```
  |                                                                                                                                                                                              
  |                                                                                                                                                                                        |   0%
```


```
  switch(datasetJson[["idType"]],
    {
     print("Checking genes in Database")
      genes_in_db <- read.csv2(paste0(workingDir,"//","storage/genes.csv"),header=TRUE,stringsAsFactors = FALSE)
      genes_in_db <- genes_in_db[genes_in_db$SPECIES==datasetJson[["species"]],]
  
      gene_id <- rep("",nrow(data_matrix))
    
      for(actGeneRow in 1:nrow(data_matrix)){
        setTxtProgressBar(pb, actGeneRow)
        
        indexFound<-c()
        if(datasetJson[["idType"]]=="ensemblid"){
          indexFound<-which(genes_in_db$ENSEMBL==rownames(data_matrix)[actGeneRow])
        }
        if(datasetJson[["idType"]]=="entrezid"){
          indexFound<-which(genes_in_db$ENTREZID==rownames(data_matrix)[actGeneRow])
        }
        if(datasetJson[["idType"]]=="symbol"){
          if(sum(genes_in_db$SYMBOL==rownames(data_matrix)[actGeneRow])==0){ #r replaces duplicated rownames in rownames with "." -> if that happend in the preprocessing of the data, we can account for this by removing the . and everything after that
            indexFound<-which(genes_in_db$SYMBOL==strsplit(rownames(data_matrix)[actGeneRow], "\\.")[[1]][1])
          }else{
            indexFound<-which(genes_in_db$SYMBOL==rownames(data_matrix)[actGeneRow])
          }
        }  
        
        indexFound<-indexFound[!is.na(indexFound)]
        
        if(length(indexFound)>0){
          gene_id[actGeneRow]<-paste0(genes_in_db$ENSEMBL[indexFound[1]],"_",genes_in_db$ENTREZID[indexFound[1]])
          foundGeneInDatabase[actGeneRow]<-length(indexFound) 
        }
      }

      datasetJson[["idType"]]<-"geneID"
      
        print(paste0("Found ",sum(foundGeneInDatabase>0)," genes (",sum(foundGeneInDatabase>1)," double) out of ",nrow(data_matrix)))
    print(paste0("Out of ",sum(foundGeneInDatabase>0)," found, ",sum(!duplicated(gene_id) & foundGeneInDatabase>0)," are not duplicates"))
    },
      "peakname"={
        print("Checking genes in Database")
        genes_in_db <- read.csv2(paste0(workingDir,"//","storage/peaks.csv"),header=TRUE,stringsAsFactors = FALSE)
    

        gene_id <- rep("",nrow(data_matrix))
      
        for(actGeneRow in 1:nrow(data_matrix)){
          setTxtProgressBar(pb, actGeneRow)
  
          indexFound<-which(genes_in_db$name==rownames(data_matrix)[actGeneRow])
          indexFound<-indexFound[!is.na(indexFound)]
          
          if(length(indexFound)>0){
            gene_id[actGeneRow]<-paste0(genes_in_db$name[indexFound[1]])
            foundGeneInDatabase[actGeneRow]<-length(indexFound) 
          }
        }
  
        
        print(paste0("Found ",sum(foundGeneInDatabase>0)," peaks (",sum(foundGeneInDatabase>1)," double) out of ",nrow(data_matrix)))
        print(paste0("Out of ",sum(foundGeneInDatabase>0)," found, ",sum(!duplicated(gene_id) & foundGeneInDatabase>0)," are not duplicates"))
      
    }
    )
```


```
[1] "Checking genes in Database"

  |                                                                                                                                                                                              
  |==============================================                                                                                                                                          |  25%
  |                                                                                                                                                                                              
  |============================================================================================                                                                                            |  50%
  |                                                                                                                                                                                              
  |==========================================================================================================================================                                              |  75%
  |                                                                                                                                                                                              
  |========================================================================================================================================================================================| 100%[1] "Found 4 genes (0 double) out of 4"
[1] "Out of 4 found, 4 are not duplicates"
```


```
   close(pb)
```


```
  count_matrix<-data_matrix[!duplicated(gene_id) & foundGeneInDatabase>0,!is.na(sample_information$region_id)]
  gene_id_matrix<-matrix(gene_id[!duplicated(gene_id) & foundGeneInDatabase>0],ncol=1)
  colnames(gene_id_matrix)<-c("gene_id")
  
  write.csv2(gene_id_matrix,paste0(outputData,"/gene_information.csv"),row.names=FALSE)
  
  
  print("Create pre aggregated count matrix and import data")
```


```
[1] "Create pre aggregated count matrix and import data"
```


```
  sample_information_filtered_aggregated<-data.frame(table(sample_information_filtered[,-1]))
  sample_information_filtered_aggregated<-sample_information_filtered_aggregated[sample_information_filtered_aggregated$Freq>0,]
  sample_information_filtered_aggregated<-cbind(paste0(1:nrow(sample_information_filtered_aggregated)),sample_information_filtered_aggregated)
  colnames(sample_information_filtered_aggregated)[1]<-"sampleID"
  colnames(sample_information_filtered_aggregated)[ncol(sample_information_filtered_aggregated)]<-"sampleCount"
  count_matrix_aggregated<-matrix(0,nrow=nrow(count_matrix),ncol=nrow(sample_information_filtered_aggregated))
  
    
  print(paste0("Aggregated sample_information to ",nrow(sample_information_filtered_aggregated)," unique combinations with an average frequency of ",mean(sample_information_filtered_aggregated$sampleCount)))
```


```
[1] "Aggregated sample_information to 9 unique combinations with an average frequency of 1.11111111111111"
```


```
  pb <- txtProgressBar(min=0, max=nrow(sample_information_filtered_aggregated), initial=0,style=3)
```


```
  |                                                                                                                                                                                              
  |                                                                                                                                                                                        |   0%
```


```
  for(actAggregatedSample in 1:nrow(sample_information_filtered_aggregated)){
    setTxtProgressBar(pb, actAggregatedSample)
    
    sampleIndizesToBeAggregated<-apply(sapply(2:ncol(sample_information_filtered),function(x){
      sample_information_filtered[,x]==sample_information_filtered_aggregated[actAggregatedSample,x]
    }),1,function(y){
      sum(y)==length(y)
    })
    
    if(sum(sampleIndizesToBeAggregated)==1){
      count_matrix_aggregated[,actAggregatedSample]<-count_matrix[,sampleIndizesToBeAggregated]
    }else{
      count_matrix_aggregated[,actAggregatedSample]<-rowSums(count_matrix[,sampleIndizesToBeAggregated])
    }
    
    
  }
```


```
  |                                                                                                                                                                                              
  |====================                                                                                                                                                                    |  11%
  |                                                                                                                                                                                              
  |=========================================                                                                                                                                               |  22%
  |                                                                                                                                                                                              
  |=============================================================                                                                                                                           |  33%
  |                                                                                                                                                                                              
  |==================================================================================                                                                                                      |  44%
  |                                                                                                                                                                                              
  |======================================================================================================                                                                                  |  56%
  |                                                                                                                                                                                              
  |===========================================================================================================================                                                             |  67%
  |                                                                                                                                                                                              
  |===============================================================================================================================================                                         |  78%
  |                                                                                                                                                                                              
  |====================================================================================================================================================================                    |  89%
  |                                                                                                                                                                                              
  |========================================================================================================================================================================================| 100%
```


```
  close(pb)
```


```
  rownames(count_matrix_aggregated)<-rownames(count_matrix)
  colnames(count_matrix_aggregated)<-sample_information_filtered_aggregated[,1]

  print("Write data")
```


```
[1] "Write data"
```


```
  options(scipen=20)
  
  write.csv2(sample_information_filtered_aggregated,paste0(outputData,"/sample_information_aggregated.csv"),row.names = FALSE)
  write.table(count_matrix_aggregated,paste0(outputData,"/count_matrix_aggregated.csv"),row.names=FALSE,col.names=FALSE,dec=".",sep=";")
                               
                    
  write.table(as.matrix(count_matrix),paste0(outputData,"/count_matrix.csv"),row.names=FALSE,col.names=FALSE,dec=".",sep=";")
  
  datasetJson[["maximumValue"]]<-paste0(max(count_matrix,na.rm=TRUE))
```


```
write(toJSON(datasetJson),outputJson)
print(paste0("Dataset ",datasetJson[["name"]]," done!"))
```


```
[1] "Dataset GeneExpressionTestdata done!"
```


```
if(computeSeurat==TRUE){
print("Compute TSNE....")
  set.seed(1899)
  
  mydata <- CreateSeuratObject(counts = count_matrix, project = datasetJson[["name"]])
  
  mydata <- FindVariableFeatures(mydata)
  mydata <- ScaleData(object = mydata, features = VariableFeatures(object = mydata))
  
  mydata <- RunPCA(
    object = mydata, features = VariableFeatures(object = mydata), verbose = F, 
    npcs = 20
  )
  
  tsne<-RunTSNE(
    object = mydata, dims = 1:10, do.fast = TRUE, check_duplicates = FALSE,
    num_threads = 10
  )
  reductions<-tsne@reductions
  save(reductions, file = paste0(outputData,"/reductions.RData"))
  save(tsne, file = paste0(outputData,"/seuratObject.RData"))
}
```


```
  print("Zip data...")
```


```
[1] "Zip data..."
```


```
  setwd(outputDir)
 
    zip(paste0(outputDir,"/",datasetJson[["name"]],"_count_matrix.zip"),flags="-q",c(gsub(outputDir,"",outputJson),
                                                                                     gsub(outputDir,"",paste0(outputData,"/coordinates_to_region_index.csv")),
                                                                                     gsub(outputDir,"",paste0(outputData,"/gene_information.csv")),
                                                                                     gsub(outputDir,"",paste0(outputData,"/sample_information_aggregated.csv")),
                                                                                     gsub(outputDir,"",paste0(outputData,"/count_matrix_aggregated.csv")),
                                                                                     gsub(outputDir,"",paste0(outputData,"/sample_information.csv")),
                                                                                     gsub(outputDir,"",paste0(outputData,"/count_matrix.csv"))))
    if(computeSeurat){
      zip(paste0(outputDir,"/",datasetJson[["name"]],"_seuratData.zip"),flags="-q",c(gsub(outputDir,"",paste0(outputData,"/coordinates_to_region_index.csv")),
                                                                                     gsub(outputDir,"",paste0(outputData,"/gene_information.csv")),
                                                                                     gsub(outputDir,"",paste0(outputData,"/sample_information_aggregated.csv")),
                                                                                     gsub(outputDir,"",paste0(outputData,"/count_matrix_aggregated.csv")),
                                                                                     gsub(outputDir,"",paste0(outputData,"/sample_information.csv")),
                                                                                     gsub(outputDir,"",paste0(outputData,"/reductions.RData")),
                                                                                     gsub(outputDir,"",paste0(outputData,"/seuratObject.RData"))))
    }
  

  unlink(outputJson, recursive = TRUE)
  unlink(outputData, recursive = TRUE)
```


```
print("Done!")
```


```
[1] "Done!"
```


```
sink()
```
