## Supplementary Data 2 for "BrainTACO: An Explorable Multi-Scale Multi-Modal Brain Transcriptomic And Connectivity Data Resource": dataset_comparison_correlation_computation.nb.html

R Notebook


Code 

- Show All Code
- Hide All Code
- Download Rmd

### R Notebook


```
library(openxlsx)
setwd(dirname(rstudioapi::getActiveDocumentContext()$path))
```


```
excelIndexToCorMatrix<-function(excelIndex){
  letter2number <- function(x) {utf8ToInt(tolower(x)) - utf8ToInt("a") + 1L}
  
  #split number from 
  for(actIndex in length(excelIndex):1){
    if(suppressWarnings(is.na(as.numeric(substr(excelIndex,actIndex,nchar(excelIndex)))))){
      return(c(letter2number(substr(excelIndex,1,actIndex)),as.numeric(substr(excelIndex,actIndex+1,nchar(excelIndex)))))
    }
  }
}

excelIndexToCorMatrixWithOffset<-function(excelIndex,excelOffset){
  excelIndexToCorMatrix(excelIndex)-(excelIndexToCorMatrix(excelOffset)-1)
}

sameCellTypesDifferenceToNotSameDatasets<-function(sameCellTypeExcelIndices,correlationResults){
  sameCellTypeIndices<-t(sapply(sameCellTypeExcelIndices,function(x){
  excelIndexToCorMatrixWithOffset(x,"C3")
  }))[,c(2,1)] #row,column format

  datasetnames<-sapply(rownames(correlationResults),function(x){
    strsplit(x,".",fixed=TRUE)[[1]][2]
  })
  notSameDatasetAndCelltypeIndices<-c()
  for(i in 1:nrow(correlationResults)){
    for(j in 1:ncol(correlationResults)){
      if(datasetnames[i]!=datasetnames[j]){
        isAlreadyInSameCellTypeIndices<-FALSE
        for(k in 1:nrow(sameCellTypeIndices)){
          if(i==sameCellTypeIndices[k,1] && j==sameCellTypeIndices[k,2]){
            isAlreadyInSameCellTypeIndices<-TRUE
          }
        }
        if(isAlreadyInSameCellTypeIndices==FALSE){
          notSameDatasetAndCelltypeIndices<-rbind(notSameDatasetAndCelltypeIndices,c(i,j))
        }
      }
    }
  }
  
  return(wilcox.test(correlationResults[sameCellTypeIndices],correlationResults[notSameDatasetAndCelltypeIndices],alternative="greater"))
}
```


```
queryResults<-read.xlsx("dataset_comparison_query_results.xlsx",sheet="Mouse_PFC")
```


```
queryResults<-read.xlsx("dataset_comparison_query_results.xlsx",sheet="Mouse_Visual_Areas")

queryResults<-queryResults[,7:ncol(queryResults)]
for(i in 1:ncol(queryResults)){
  queryResults[,i]<-as.numeric(queryResults[,i])
}


correlationResults<-cor(queryResults,method = "spearman",use = "pairwise.complete.obs")

write.xlsx(as.data.frame(correlationResults), file = "query_results_correlation_single_xlsx/Mouse_Visual_Areas.xlsx",overwrite=TRUE)

sameCellTypeExcelIndices<-c("I3","J3","K4","L5","M6","N7","O8","J9","I10","C9","C10","D11","E12","F13","G14","H15")
sameCellTypesDifferenceToNotSameDatasets(sameCellTypeExcelIndices,correlationResults)
```


```
    Wilcoxon rank sum test with continuity correction

data:  correlationResults[sameCellTypeIndices] and correlationResults[notSameDatasetAndCelltypeIndices]
W = 696, p-value = 0.2926
alternative hypothesis: true location shift is greater than 0
```


```
queryResults<-read.xlsx("dataset_comparison_query_results.xlsx",sheet="Mouse_Striatum")

queryResults<-queryResults[,7:ncol(queryResults)]
for(i in 1:ncol(queryResults)){
  queryResults[,i]<-as.numeric(queryResults[,i])
}


correlationResults<-cor(queryResults,method = "spearman",use = "pairwise.complete.obs")

write.xlsx(as.data.frame(correlationResults), file = "query_results_correlation_single_xlsx/Mouse_Striatum.xlsx",overwrite=TRUE)

sameCellTypeExcelIndices<-c("C11","C12","D13","E14","F15","G16","H17","I18","J19","L11","K3","L3","M4","N5","O6","P7","Q8","R9","S10")
sameCellTypesDifferenceToNotSameDatasets(sameCellTypeExcelIndices,correlationResults)
```


```
    Wilcoxon rank sum test with continuity correction

data:  correlationResults[sameCellTypeIndices] and correlationResults[notSameDatasetAndCelltypeIndices]
W = 2288.5, p-value = 2.818e-07
alternative hypothesis: true location shift is greater than 0
```


```
queryResults<-read.xlsx("dataset_comparison_query_results.xlsx",sheet="Mouse_Hypothalamus")

queryResults<-queryResults[,7:ncol(queryResults)]
for(i in 1:ncol(queryResults)){
  queryResults[,i]<-as.numeric(queryResults[,i])
}


correlationResults<-cor(queryResults,method = "spearman",use = "pairwise.complete.obs")

write.xlsx(as.data.frame(correlationResults), file = "query_results_correlation_single_xlsx/Mouse_Hypothalamus.xlsx",overwrite=TRUE)

sameCellTypeExcelIndices<-c("J3","K3","L4","M5","O7","P8","Q9","S3","T4","U5","V6","X8","Y9","S10","S11","K10","C10","C11","D12","E13","G15","H16","I17","T12","U13","W14","X16","Y17","Z18","C19","D20","E21","F22","H24","I25","J19","K19","L20","M21","N23","P24","Q25","R26")
sameCellTypesDifferenceToNotSameDatasets(sameCellTypeExcelIndices,correlationResults)
```


```
    Wilcoxon rank sum test with continuity correction

data:  correlationResults[sameCellTypeIndices] and correlationResults[notSameDatasetAndCelltypeIndices]
W = 12728, p-value = 4.266e-13
alternative hypothesis: true location shift is greater than 0
```


```
queryResults<-read.xlsx("dataset_comparison_query_results.xlsx",sheet="Mouse_Hippocampus")

queryResults<-queryResults[,7:ncol(queryResults)]
for(i in 1:ncol(queryResults)){
  queryResults[,i]<-as.numeric(queryResults[,i])
}


correlationResults<-cor(queryResults,method = "spearman",use = "pairwise.complete.obs")

write.xlsx(as.data.frame(correlationResults), file = "query_results_correlation_single_xlsx/Mouse_Hippocampus.xlsx",overwrite=TRUE)

sameCellTypeExcelIndices<-c("C4","D3","G3","G4","H5","I6","F9","E8","D7","C7")
sameCellTypesDifferenceToNotSameDatasets(sameCellTypeExcelIndices,correlationResults)
```


```
Warning in wilcox.test.default(correlationResults[sameCellTypeIndices],  :
  cannot compute exact p-value with ties
```


```
    Wilcoxon rank sum test with continuity correction

data:  correlationResults[sameCellTypeIndices] and correlationResults[notSameDatasetAndCelltypeIndices]
W = 152, p-value = 0.01162
alternative hypothesis: true location shift is greater than 0
```


```
queryResults<-read.xlsx("dataset_comparison_query_results.xlsx",sheet="Human_Frontal_Cortex")

queryResults<-queryResults[,7:ncol(queryResults)]
for(i in 1:ncol(queryResults)){
  queryResults[,i]<-as.numeric(queryResults[,i])
}


correlationResults<-cor(queryResults,method = "spearman",use = "pairwise.complete.obs")

write.xlsx(as.data.frame(correlationResults), file = "query_results_correlation_single_xlsx/Human_Frontal_Cortex.xlsx",overwrite=TRUE)

sameCellTypeExcelIndices<-c("D3","E3","C4","C5","D5","E4","O3","O4","O5","P6","Q7","R8","S9","T10","U11","V12","W13","X14","C15","D15","E15","F16","G17","H18","I19","J20","K21","L22","M23","N24")
sameCellTypesDifferenceToNotSameDatasets(sameCellTypeExcelIndices,correlationResults)
```


```
    Wilcoxon rank sum test with continuity correction

data:  correlationResults[sameCellTypeIndices] and correlationResults[notSameDatasetAndCelltypeIndices]
W = 5408, p-value = 5.799e-05
alternative hypothesis: true location shift is greater than 0
```


```
queryResults<-read.xlsx("dataset_comparison_query_results.xlsx",sheet="Human_Visual_Cortex")

queryResults<-queryResults[,7:ncol(queryResults)]
for(i in 1:ncol(queryResults)){
  queryResults[,i]<-as.numeric(queryResults[,i])
}


correlationResults<-cor(queryResults,method = "spearman",use = "pairwise.complete.obs")

write.xlsx(as.data.frame(correlationResults), file = "query_results_correlation_single_xlsx/Human_Visual_Cortex.xlsx",overwrite=TRUE)

sameCellTypeExcelIndices<-c("D3","E3","E4","C4","C5","D5","N3","N4","N5","O6","P7","Q8","R9","S10","T11","U12","V13","M22","L21","K20","J19","I18","H17","G16","F15","E14","D14","C14")
sameCellTypesDifferenceToNotSameDatasets(sameCellTypeExcelIndices,correlationResults)
```


```
    Wilcoxon rank sum test with continuity correction

data:  correlationResults[sameCellTypeIndices] and correlationResults[notSameDatasetAndCelltypeIndices]
W = 4092, p-value = 0.0002527
alternative hypothesis: true location shift is greater than 0
```
